# A toolkit for programmable transcriptional engineering across eukaryotic kingdoms

**DOI:** 10.64898/2026.02.10.705154

**Authors:** Izaiah J. Ornelas, Lauren A. Owens, Simon Alamos, Niklas F.C. Hummel, Mitzi G. Hernández Zamora, Ngan T. Phan, Rithu K. Pattali, Patrick M. Shih, James K. Nuñez

## Abstract

Chromatin is essential for eukaryotic life. Tens of thousands of chromatin regulator (CR) proteins exist in eukaryotic genomes that are predicted to modulate chromatin states; however, their molecular functions remain largely untested experimentally. Here, we construct a library of over 300 full-length CRs from humans, plants, yeast, protozoa, and virus, each fused to DNA-binding domains, and test their direct effect on transcriptional repression and activation in plants and human cells. We discover CRs with cross-kingdom functionality when transferred across eukaryotes, including CRs that outperform existing tools for programmable transcriptional repression and activation in plants and human cells. Using pooled CRISPR screens, we demonstrate a suite of CRISPR repressors that titrate gene expression at intermediate levels. Finally, we identify RCOR1 and MTA2 as universal eukaryotic repressors that retain repressive activity in plants, yeast, and human cells. Our toolkit advances synthetic eukaryotic engineering and expands our understanding of CR functionality across eukaryotes.

## INTRODUCTION

Eukaryotes package their genomes into chromatin that lays the foundation for DNA-templated processes such as transcription, repair, and genome organization^1,2^. Chromatin is dynamic in nature, reversibly switching between relaxed euchromatin and compacted heterochromatin.

Proteins broadly referred to as chromatin regulators (CRs) can modulate chromatin states by reading, writing, and erasing post-translational epigenetic modifications or active remodeling of chromatin structure^3–7^. Along with transcription factors (TFs), CRs are key players in regulating eukaryotic gene expression and genome organization^8^. Recent phylogenetic reconstruction and proteomics of chromatin across highly divergent eukaryotes shows high conservation of CRs and histone modifications, highlighting the selective pressure for chromatin regulation throughout eukaryotic evolution^9^. Although genomics and proteomics approaches have identified the presence of CRs in all eukaryotes, the ability to experimentally test the functional activities of CRs within and across diverse eukaryotic species remains a challenge.

Given the wide range of applications for CRs in genome engineering and synthetic biology, there is great interest in expanding the toolkit to control chromatin state and gene expression in eukaryotic cells^10–12^. Typically, CRs can be recruited to or directly fused to a programmable DNA binding domain (DBD) protein that targets the promoter of a gene, thereby enabling direct measurement of gene expression changes induced by CR recruitment^13–16^. For example, over two hundred full-length CRs from the yeast genome were synthetically fused to a zinc finger DBD and tested for transcriptional modulation of a synthetic gene reporter, ultimately yielding tools for site-specific repression and activation in *S. cerevisiae*^17^. Moreover, recent technological advances have enabled functional testing of thousands of CRs derived from human and viral genomes in a pooled or arrayed format in human cell lines^18–23^. These approaches rely frequently on amino acid fragments rather than full-length CRs. Many CRs possess multiple protein domains for their functional activity and thus, testing fragments offers partial or incomplete views of CR function^24–26^. In general, functional testing of full-length CRs encoded in different eukaryotic genomes remains less common due to the cost barriers associated with synthesis of larger, whole-protein constructs, leaving major gaps in our understanding of how CR functionality is conserved or divergent across eukaryotic species.

Most synthetic chromatin engineering efforts have focused on well-established model systems such as yeast and human cell lines. However, there is growing interest in extending these approaches to less-studied eukaryotic systems. For instance, precise transcriptional control in plants holds promise for improving agronomic traits, yet most genome engineering tools in plants are derived from studies in human cells and yeast rather than plant systems^27–29^. Beyond plants, there exist six major eukaryotic supergroups that span over a billion years of evolutionary divergence, encompassing a vast range of unique biology and chemistries^30^. Despite this diversity, it is thought that many major chromatin modifications are highly conserved across eukaryotes^31,32^. Nonetheless, our ability to manipulate non-model eukaryotes remains limited, and it is not clear how CRs from one species may function in another. The lack of cross-species engineering systems raises the need for tools that are both versatile and broadly compatible across different host organisms^33^. Rather than needing to screen for functional CRs in each new species, developing CRs that are readily deployable across diverse eukaryotic hosts will accelerate both fundamental research and applied efforts – from disease-relevant parasites (e.g., Apicomplexans), to crops (e.g., plants and algae), to organisms with biotechnological potential (e.g., filamentous fungi and diatoms).

Here, we present a large-scale platform to dissect the functional roles of hundreds of full-length CRs that originate from the genomes of humans, plants, yeast, a parasitic protozoon, and giant viruses in gene regulation. To assess conservation of function across species, we tested each CR for its ability to modulate gene expression in the model plant *Nicotiana benthamiana,* human cells, and, for a subset, in the budding yeast *Saccharomyces cerevisiae* to identify CRs that retain repressive activity across host backgrounds. Together, we discover new programmable CRs that outperform current state-of-the-art technologies for transcriptional control in humans and plants and uncover repressors with conserved activity across biological kingdoms, offering a general toolkit for modulating gene regulation that can be applied broadly across biology, from basic research to translational applications.

## RESULTS

### A multi-kingdom CR library enables programmable gene modulation across species

To enable systematic discovery of chromatin regulator (CR) activities across eukaryotic genomes, we constructed a library of full-length CR genes derived from diverse phylogenetic sources and with varying predicted molecular functions (**Figure 1A** and Table S1). We reasoned that using evolutionary diverse CRs could reveal conserved and context-dependent mechanisms of gene regulation while also providing numerous effectors for engineering programmable transcriptional modulation. To this end, we curated a library of 423 CR genes from: *Arabidopsis thaliana* (n = 136), *Homo sapiens* (n = 95), *Sorghum bicolor* (n = 86), *Saccharomyces cerevisiae* (n = 60), *Toxoplasma gondii* (n = 35), and *Megaviridae* (n = 11) (**Figure 1B**). Including CRs from the Apicomplexan parasite *Toxoplasma gondii* and giant viruses extends our library beyond the canonical eukaryotic supergroups and broadens the search for CRs that may function across different hosts^34–36^.

**Figure 1.**
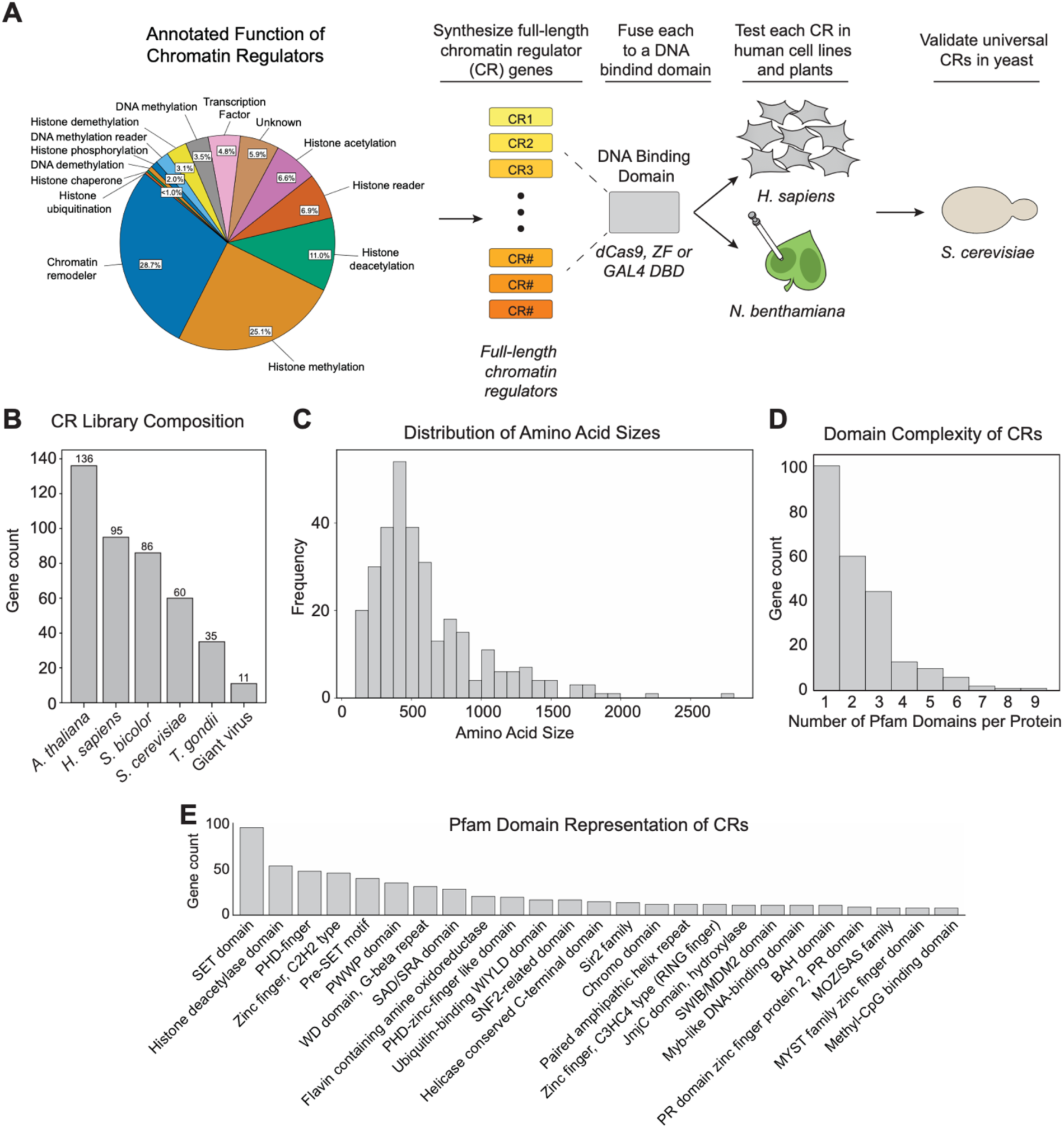
Design and composition of the Chromatin Regulator (CR) library. **(A)** Functional annotation of CRs in the library, pie chart depicts the distribution of annotated functions. Schematic (right) illustrates cloning of full-length CRs as programmable DNA-binding fusions (dCas9, ZF, or GAL4 DBD) for testing across species (*H. sapiens*, *N. benthamiana*, and *S. cerevisiae*). **(B)** Bar plot showing species composition of the CR library per species. **(C)** Distribution of CR amino acid sizes in the library. **(D)** Pfam domain representation of CRs. Bar plots show the top 25 most frequent domains identified across the library. **(E)** Domain complexity of CRs, histogram shows the number of annotated Pfam domains per protein.

Each CR in the library was designed for direct fusion to modular DBDs, such as catalytically dead Cas9 (dCas9), synthetic zinc fingers, or the GAL4 DBD. These fusions enable us to recruit CRs to the promoters of either endogenous genes or synthetic reporters, establishing a toolkit for programmable gene regulation across multiple organisms. Importantly, CRs in our library vary in amino acid length, with proteins ranging from ∼200 to over 2,500 amino acids and an average frequency around 500 residues (**Figure 1C**). This size variation reflects the complexity of CRs, many of which are large, multidomain proteins with scaffolding or enzymatic activities that are essential for their function in chromatin biology. While previous protein fragment-based approaches have been powerful for identifying minimal effector domains^18–21^, our approach of using full-length proteins preserves their native structure and function.

To further characterize the molecular diversity of our library, we analyzed Pfam domain content across all full-length CR sequences. We observe broad representation of canonical chromatin regulatory domains, including SET domains, histone deacetylase domains, PHD fingers, chromodomains, BAH domains, and various zinc finger motifs (**Figure 1D**). Many CRs contain multiple annotated Pfam domains per protein, with some proteins having as many as 8-9 annotated domains (**Figure 1E**), reflecting the structural complexity needed to perform their diverse regulatory functions and mediate interactions at specific genomic regions.

With the library in hand, we sought to experimentally test each CR in two highly divergent eukaryotic kingdoms: Plantae and Animalia. We posit that heterologous expression of chromatin regulators in distinct evolutionary lineages enables testing of their functional conservation. In addition, this approach allows discovery of CRs whose activities could be masked by endogenous regulation, potentially improving or altering their function in a heterologous cellular background. For example, the human DNA demethylase TET1 has been utilized successfully in plants to alter DNA methylation and transcriptional regulation, demonstrating how cross-kingdom transfer of CRs can be powerful in organisms with limited tools ^27^.

Taken together, our multi-kingdom, full-length CR library establishes a versatile platform for programmable gene modulation across species, enabling high-throughput and locus-specific investigations into CR based mechanisms of transcriptional regulation and providing a platform for uncovering both conserved and context specific modes of epigenetic gene regulation.

### Characterization of CRs *in planta*

Compared to the extensive tool development in yeast and human systems, plant genetic engineering generally relies on a limited set of trans-elements largely borrowed from yeast and humans. While cis-regulatory elements such as promoters and enhancers have been leveraged across plant systems for gene modulation, trans-elements like targeted recruitment of CRs remain less characterized^34–36^. Only a handful of trans-elements have been characterized for plant genetic engineering, with many of them derived from previous yeast or human cell studies (e.g., the herpes simplex virus-derived transcriptional activator VP16)^37,38^. While recent efforts have begun to identify and use endogenous plant proteins as engineering tools, the broader toolkit remains narrow ^39–41^. Our CR library provides us with the opportunity to expand the plant engineering toolkit by testing CRs from highly divergent lineages to discover new regulators that function in plants.

We tested the CR library using an agroinfiltrated synthetic transcriptional system previously developed for assaying transcriptional activators and repressors in *Nicotiana benthamiana*^25^ (**Figures 2A-B**).Transient expression in *N. benthamiana* leaves can generate data in a few days, compared to the months-long process of generating stable transgenic plant lines, making it an ideal method for screening the activity of large protein libraries^42^. Our synthetic system consisted of N-terminal fusions of each CR to a GAL4 DBD and a GFP reporter construct. When co-expressed with the fluorescent reporter, the GAL4 DBD targeted the CR to binding sites upstream the promoter region of the reporter, enabling the CR to modulate GFP expression. To measure the dynamic range of the system, we used GAL4 DBD-VP16 as a control for activators^43^ and Gal4 DBD-SUPERMAN as a control for repressors^44^. CR activity on GFP expression was normalized to our neutral control, GAL4 DBD without a CR (GAL4 DBD alone) co-expressed with the reporter. CRs with a fold change GFP greater than 2.0 were classified as activators, and CRs with a fold change less than 0.7 were classified as repressors. CRs with activity in between these cutoffs were considered neutral.

**Figure 2.**
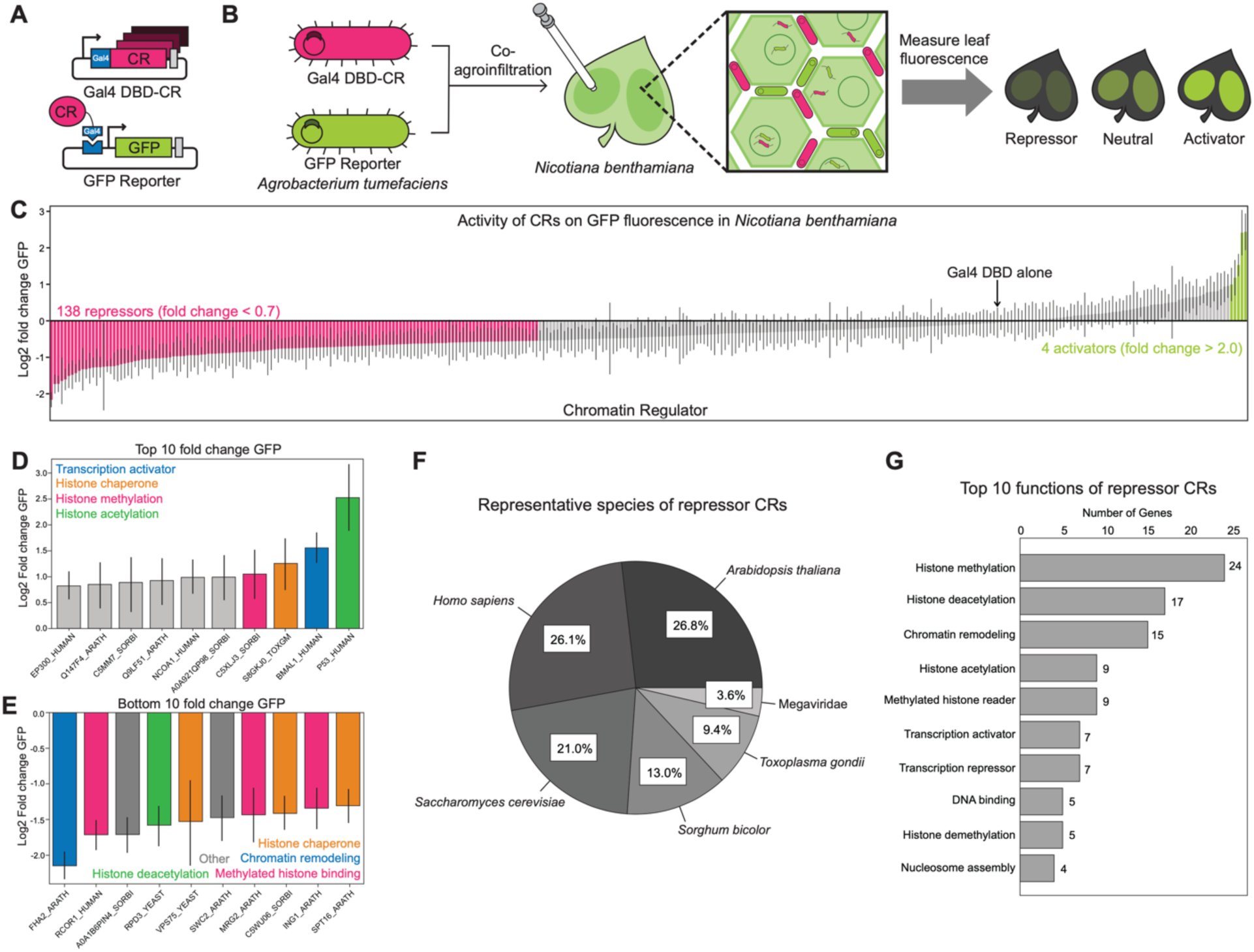
Characterization of CRs in planta identifies effective repressors and activators. **(A)** Schematic of the plant reporter assay, full-length CRs were fused to the GAL4 DBD and co-expressed with a GFP reporter plasmid in *A. tumefaciens*. **(B)** Experimental workflow of co-agroinfiltration into *N. benthamiana* leaves. CR fusions were delivered alongside the GFP reporter, and fluorescence was used to quantify repressive, neutral, or activity effects on the reporter. **(C)** Activity of CRs on GFP reporter. Log₂ fold-change GFP values are shown relative to GAL4 DBD alone (indicated by the black arrow). Bars represent individual CRs, with repressors (pink), neutral (gray), and activators (green) distributed across the library. Error bars represent the standard deviation. **(D)** Bar plot showing the top 10 log₂ fold-change GFP CRs identified from plant screen. Light gray CRs were not activators. Error bars represent the standard deviation. **(E)** Bar plot showing the bottom 10 log₂ fold-change GFP CRs identified from plant screen. **(F)** Species distribution of CRs identified as repressors in *N. benthamiana*. Error bars represent the standard deviation. **(G)** Top annotated functions of repressor CRs in the assay. The bar chart shows the most frequent functional categories.

Of the CRs tested in *N. benthamiana*, 197 were neutral, 138 were repressors, and 4 were activators (**Figure 2C**). Of the four CRs that acted as activators in *N. benthamiana*, two were *H. sapiens* genes (P53_HUMAN and BMAL1_HUMAN), one was a *S. bicolor* gene (C5XLJ3_SORBI), and one was a *T. gondii* gene (S8GKJ0_TOXGM) (**Figure 2D**). P53_HUMAN and BMAL1_HUMAN are annotated as transcription factors, C5XLJ3_SORBI is annotated as a histone lysine methyltransferase, and the *T. gondii* gene is annotated as an ASF1-like histone chaperone. The strongest of these, P53_HUMAN (mean fold change GFP = 5.76), a master transcriptional regulator and important gene in cancer regulation, increased GFP expression levels on par with VP16 (mean fold change GFP = 5.72)^45^. Plants do not have a homolog to P53_HUMAN, but a previous study identified increased homologous recombination rates and early senescence and fasciation in transgenic *A. thaliana* expressing P53_HUMAN, implicating the ability of P53_HUMAN to modulate plant gene expression^46^. P53_HUMAN contains two trans activation domains (TADs), but its ability to activate gene expression had not been tested *in planta*^47^.

The 138 CRs that we identified as repressors in *N. benthamiana* were diverse in their origin and predicted functions. We found CRs from each sampled organism with repressive activity (*A. thaliana* = 37, *H. sapiens* = 36, *S. cerevisiae* = 29, *S. bicolor* = 18, *T. gondii* = 13, Megaviridae = 5) (**Figure 2F**) and a variety of annotated functions. The five most frequent annotated mechanisms of the repressor CRs are histone methylation (n = 24), histone deacetylation (n = 17), chromatin remodeling (n = 11), histone acetylation (n = 9), and transcription repression (n = 7) (**Figure 2G**). The strongest repressors from the plant screen originated from multiple species and had annotated functions related to known chromatin silencing mechanisms (**Figure 2E**). FHA2_ARATH, a plant-specific forkhead-domain subunit of the *Arabidopsis thaliana* ISW1 chromatin remodeling complex, was the strongest repressor in *N. benthamiana* (fold change GFP = 0.23), comparable to the state-of-the-art repressive activity of the EAR domain motif SRDX in some contexts^48^. The *S. bicolor* homolog of FHA2_ARATH, A0A1B6PIN4_SORBI, was also one of the strongest repressors (fold change GFP = 0.31). A subunit of the *H. sapiens* BHC histone demethylase and deacetylase complex, RCOR1_HUMAN (fold change GFP = 0.30), the *S. cerevisiae* histone deacetylase RPD3_YEAST (fold change GFP = 0.33), and the *S. cerevisiae* histone chaperone VPS75_YEAST (fold change GFP = 0.35) also induced notable repressive activity on reporter expression^49–51^. Identification of these strong repressors expands the available toolkit for tuning gene repression in plants. As they function through a variety of mechanisms such as chromatin remodeling and histone deacetylation, these tools enable context-dependent tuning of gene expression.

Additionally, we identified several CRs with unexpected activity on gene expression. For example, both catalytic and non-catalytic subunits of the human HBO1 histone acetyltransferase complex, KAT7_HUMAN, JADE1_HUMAN, and ING4_HUMAN displayed repressor activity (fold change GFP = 0.61, 0.51, and 0.47, respectively) despite annotations indicating activity typically involved in transcriptional activation. The HBO1 HAT complex deposits histone 3 lysine 14 (H3K14), H4K5, H4K8, and H4K12 acetyl marks with KAT7_HUMAN, also known as HBO1, a MYST family histone acetyltransferase, being the catalytic subunit of the complex^52,53^. While it does not have catalytic activity, JADE1_HUMAN contains two PHD zinc finger domains and acts as a scaffolding subunit^53^. JADE1_HUMAN coordinates the complex to and mediates interactions between the complex and chromatin^54^. HBO1 complexes containing JADE proteins, as opposed to BRPFs, specifically catalyze H4 acetylation. ING4_HUMAN contains a PHD domain, which enables binding to methylated and acetylated lysine residues^55^. While MYST HATs are conserved in plants, the HBO1 complex is not^56^. Histone acetylation is broadly associated with gene activation across eukaryotes. Thus, the mechanism through which the HBO1 subunits are repressing gene expression in *N. benthamiana* is unclear^57^.

Our findings highlight how heterologous expression of CRs can uncouple protein function from native context, potentially leading to unexpected regulatory outcomes. Taken together, we discover a set of repressors that control transcription through various mechanisms and at different strengths, which enables further development of a tunable toolkit for controlling transcription *in planta* (**Figure 2C and 2G**).

### Functional profiling of CR transcriptional activity in human cells

Next, we utilized the CR library to modulate the expression of endogenous genes in human cells. We cloned each CR to the C-terminus of dCas9 with a blue fluorescent protein (BFP) serving as a linker, which allows us to measure dCas9-CR expression once expressed in cells (Figure S1A). We designed an arrayed screen that allows us to measure gene expression changes induced by each dCas9-CR in two different HEK293T lines, each with an non-essential endogenous gene tagged with GFP (*CLTA-GFP* and *RAB11A-GFP*)^58^, thus allowing us to perturb two native regulatory contexts. Each cell line constitutively expresses an sgRNA targeting the promoter of the tagged gene and gene expression changes were measured quantitatively by CLTA-GFP or RAB11A-GFP protein levels using single cell flow cytometry, as shown previously^59^ (**Figure 3A**). We utilize CRISPRi-KRAB (KOX1) as a positive strong repressor^60^, dCas9 as a mild repressor, and dCas9-VP64-p65-Rta (VPR) as an activator^61^ (**Figure 3B**). We note that *CLTA* and *RAB11A* are expressed natively at high levels, thus allowing us to discover potent activators that can further increase their expression.

**Figure 3.**
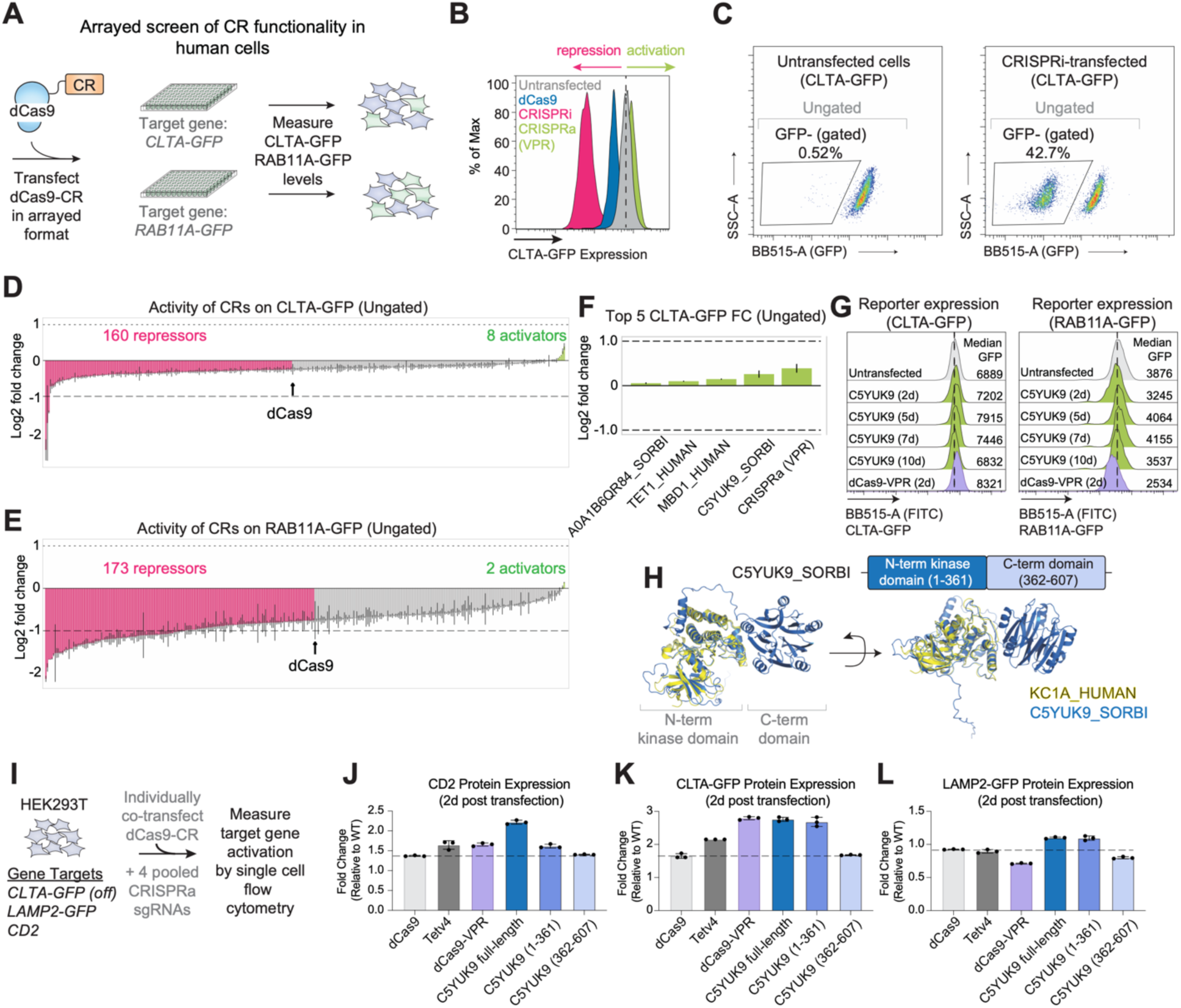
Functional profiling of CR transcriptional activity in human cells. **(A)** Schematic of the experimental workflow for transient transfection of dCas9-CR fusions in HEK293T cells with sgRNAs targeting endogenous genes tagged with GFP (CLTA-GFP and RAB11A-GFP). **(B)** Example flow cytometry histograms CLTA-GFP for untransfected, dCas9, and CRISPRi controls. **(C)** Representative flow cytometry plots showing ungated and GFP-gated populations used to measure reporter fold change. In untransfected CLTA-GFP cells, only background GFP-events are detected, whereas CRISPRi-transfected cells show a large GFP-population, confirming silencing and the gating strategy used for downstream analysis. **(D)** Activity of CRs on CLTA-GFP reporter in human cells 5 days post-transfection. Log₂ fold-change values are shown relative to untransfected control. Bars represent the mean ± s.d. of two biological replicates of CRs, with repressors (pink) and activators (green) distributed across the library. **(E)** same as in **(C)** but for RAB11A-GFP reporter in human cells 5 days post-transfection. **(F)** Top five log₂ fold-changes for CLTA-GFP from the ungated analysis. **(G)** Representative flow cytometry histograms showing the distribution of GFP expression for C5YUK9_SORBI throughout the time course at CLTA-GFP and RAB11A-GFP reporters. **(H)** Two angles of a structure alignment between the predicted structure of C5YUK9_SORBI from AlphaFold, showing the N-terminal kinase-like domain and C-terminal domain, and the human protein KC1A_HUMAN (yellow) that aligns to the kinase-like domain of C5YUK9_SORBI. **(I)** Schematic of the activation assay workflow testing dCas9-CRs for transcriptional activation across multiple targets (CD2, CLTA-GFP, LAMP2-GFP). **(J)** Quantification of target gene fold-change by flow cytometry two days post-transfection compared to untransfected cells. Data are presented as mean ± s.d. of 3 biological replicates. **(K)** Same as in **(J)** but for CRISPR-off silenced CLTA-GFP reporter in human cells 2 days post-transfection. **(L)** Same as in **(J)** but for LAMP2-GFP reporter in human cells 2 days post-transfection.

To compare CR activity systematically, we focused on Day 5 post-transfection of dCas9-CR expressing plasmids when GFP silencing and activation are near maximum and we calculated GFP fold change relative to baseline expression for each CR. We first identified functional CRs by measuring median GFP levels in transfected cells (ungated cell population, **Figure 3C**). CRs with fold change below dCas9 were classified as repressors, whereas CRs driving expression above baseline were identified as activators. Our analysis revealed a full spectrum of activity across CLTA and RAB11A reporters, spanning strong repression to measurable activation (**Figures 3D-E** and S1B-C). Using these metrics, we identified 160 repressors and 8 activators for CLTA-GFP, and 173 repressors and 2 activators for RAB11A-GFP.

### A plant protein induces robust programmable gene activation in human cells

Surprisingly, only one CR, C5YUK9_SORBI, produced consistent activation of CLTA and RAB11A at levels comparable to our positive control dCas9-VPR (**Figures 3F-G** and S1D). C5YUK9_SORBI is an uncharacterized protein with a predicted serine/threonine protein kinase domain based on AlphaFold structure prediction^62^. We used Dali to perform a search for known human proteins with similar structures, which yielded KC1A_HUMAN as the top hit^63^. The alignment is restricted primarily to the AlphaFold structure kinase domain of KC1A, whereas the C-terminal domain of C5YUK9 has low structural homology to proteins in the human proteome (**Figure 3H**).

To validate the activation activity of C5YUK9_SORBI, we next tested its ability to activate expression of multiple endogenous genes in HEK293T cells. We selected the endogenous gene *CD2* and two fluorescently tagged genes, *LAMP2-mNeonGreen* and *CLTA-GFP*, the latter of which was previously silenced epigenetically via DNA methylation and H3K9me3 using CRISPRoff^59^. We chose these targets due to their relatively low expression compared to our previous reporters, providing a broader dynamic range to assess activation strength across different genes. Cells were co-transfected with plasmids expressing dCas9-CRs and four predicted CRISPRa sgRNAs per target^64^ (**Figure 3I**). Two days post-transfection, protein levels of the targets were measured by flow cytometry in single cells and we calculated fold change relative to wild type levels. Consistent with its activity on our original reporters, C5YUK9_SORBI induced the highest CD2 expression, while unexpectedly outperforming dCas9-VPR and TETv4 that fuses dCas9 to the catalytic core of TET1 DNA demethylase^59^ (**Figure 3J**). Similarly, C5YUK9_SORBI increased CLTA-GFP and LAMP2-mNeonGreen expression, comparable to and exceeding our positive control activator (**Figures 3K-L**). Notably, the kinase domain of C5YUK9_SORBI retained most of the activation activity, suggesting its core transcriptional activation domain resides within this region. Although the mechanism of C5YUK9-mediated activation is unclear, previous work has shown that programmable phosphorylation of histone H3 at serine 28 (H3S28ph) with dCas9-MSK1 can induce transcriptional activation^65^.

Our data highlight our unexpected discovery of a non-human CR that drives robust activation of endogenous human genes at levels similar to a state-of-the-art CRISPR activation platform. The activity of C5YUK9 indicates that CRs from different species can function effectively in human cells. This suggests that heterologous proteins may bypass endogenous regulation and retain their function outside their native context, as shown previously with human TET1 for DNA demethylation in plants^27^.

### Repressive activity of the CR library in human cells

To identify the strongest CRs, we next analyzed the GFP negative (GFP-) population rather than the ungated population (**Figure 3C**). While the ungated measurement shows overall shifts in reporter expression across the entire cell population, the GFP-gate selects for cells that have measurably reduced reporter expression compared to unperturbed cells, allowing us to quantitatively rank CRs by their repression strength. We define relative activity as the proportion of GFP-negative cells normalized to transfection efficiency, measured using a BFP marker at day 2 post-transfection (Figure S1A). GFP expression was measured over a time course (2-, 5-, 7-, 10-, and 14-day post-transfection) to capture early repression and memory of silencing as cells dilute the plasmids expressing the dCas9 fusions throughout subsequent cell divisions.

Our time course analysis yielded a range of target gene expression dynamics across CRs. dCas9 alone showed minimal repression across the reporter genes, while the positive repressor control CRISPRi (dCas9-KRAB) displayed the strongest repression across all CRs tested (**Figure 4A** and S1E). Generally, CR activity peaks at 5 days post-transfection and weakens throughout the time course, highlighting that the dCas9-CRs in our library do not induce long-term epigenetic silencing of the gene reporters. Two notable exceptions are KRAB and TIF1B (TRIM28/KAP1), which are known physical interactors that establish repressive H3K9me3 and can induce longer-term epigenetic memory^66^. Of note, TRIM28 is a large protein with at least seven distinct domains that together function as a hub for epigenetic silencing^67^, highlighting the advantage of synthesizing full-length, multidomain CRs that can recapitulate their native biochemical activities in living cells.

**Figure 4.**
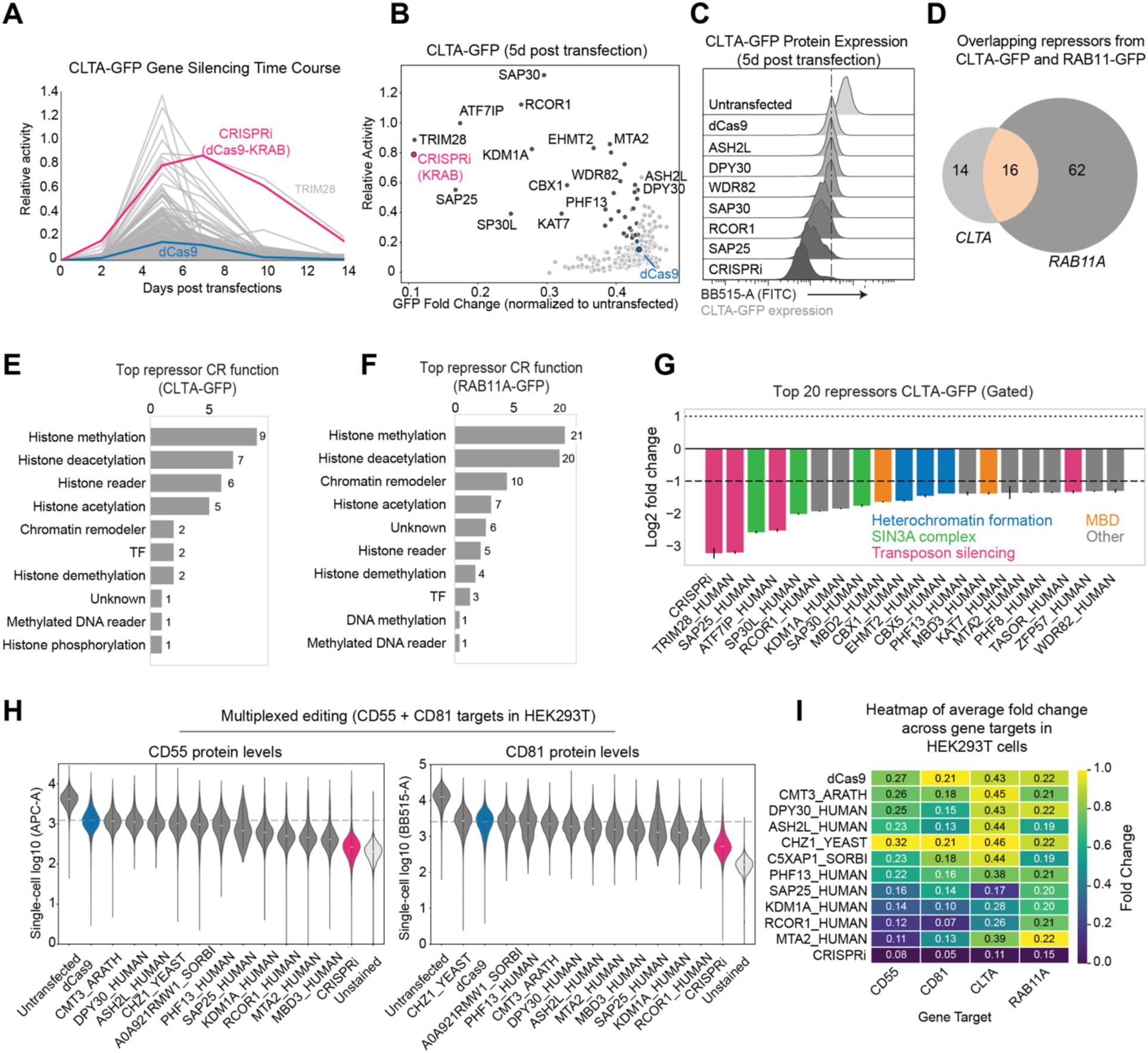
Repressive activity of the CR library in human cells. **(A)** Time course of GFP silencing for CLTA-GFP reporter across tested CRs. **(B)** Scatterplot of relative activity and GFP fold change for CLTA-GFP at 5 days post transfection. **(C)** Representative GFP distributions from flow cytometry for selected CRs in CLTA-GFP. **(D)** Overlapping repressors identified from CLTA-GFP and RAB11A-GFP reporters. **(E)** Top 10 annotated functions of repressive CRs in the assay for CLTA-GFP. The bar chart shows the most frequent functional categories. **(F)** Top 10 annotated functions of repressive CRs in the assay for RAB11A-GFP. The bar chart shows the most frequent functional categories. **(G)** Bar plot showing the CLTA-GFP log₂ fold-change for top 20 repressors identified from human screen for the gated population. Fold changes were calculated using median GFP from GFP-gate compared to untransfected control. Bars are colored by annotated function or pathway. **(H)** Violin plots showing single-cell antibody fluorescence intensity for CRs with lower median expression than dCas9 at *CD55* and *CD81*. **(I)** Heatmap summarizing per-gene min–max–normalized average GFP fold changes across reporter assays for individual CRs.

To distinguish the strongest repressor candidates, we focused on CRs that induced higher relative repressive activity with lower GFP fold-change and higher relative activity compared to dCas9 control (**Figures 4B-C** and S1F-G). We identify 92 CRs that induced stronger repressive activity compared to dCas9 alone, including 14 that are specific to CLTA-GFP, 62 specific to RAB11A-GFP and 16 shared across both reporters (**Figures 4D** and S1H). Of the 92 total repressors, about half were human and nearly all others were plant-derived, except for TTI1_YEAST, SUS1_YEAST, IES5_YEAST, and SNT1_YEAST. Functional annotation of the top repressors highlighted enrichment for canonical silencing mechanisms, with histone methylation, histone deacetylation, and chromatin remodelers as the most frequently represented (**Figures 4E-F**).

The top 20 repressors for CLTA-GFP show that CRs within the same complex or pathway induce comparable repression strength, consistent with their distinct mechanisms in chromatin-based silencing (**Figure 4G**). For example, CRISPRi (dCas9-KRAB), TIF1B_HUMAN (TRIM28), MCAF1_HUMAN (ATF7IP), and TASOR_HUMAN are involved in transposon silencing and were among the strongest repressors for CLTA^68,69^. Members of the SIN3A complex, including SAP25_HUMAN, SP30L_HUMAN, and SAP30_HUMAN, which recruit the SIN3A-HDAC complex, were in the top eight repressors^70,71^. Similarly, proteins involved in heterochromatin formation, CBX1_HUMAN (HP1-β), CBX5_HUMAN (HP1-α), and EHMT2_HUMAN (G9A), induced similar strengths of repression^72,73^. Lastly, MBD2_HUMAN and MBD3_HUMAN, which read DNA methylation and recruit additional repressive factors, produced repression at similar levels^74^.

Surprisingly, RAB11A repression is more permissive to non-human CRs. Among the top 20 repressors (Figure S1I), numerous plant-derived CRs showed activity alongside canonical human repressors. These included A0A1I9LNM3_ARATH, a histone methyltransferase, and C5XAP1_SORBI, a SET-domain-containing protein, both of which likely mediate repression through histone methylation. Additionally, HDA9_ARATH, a histone deacetylase known to regulate gene silencing in plants, repressed RAB11A-GFP^75^. These findings show that RAB11A can be effectively repressed by CRs originating from different species.

Interestingly, WDR82_HUMAN, ASH2L_HUMAN, and DPY30_HUMAN, all canonical members of the Set1/COMPASS complex that deposits transcriptional activating H3K4 trimethylation, induced reproducible repression although at weaker levels compared to CRISPRi^76^. This activity points to the possible context dependent interactions of CRs across different genes as shown previously in mammalian cells^77,78^. Together, our screen in human cells revealed a broad range of CR activity, highlighting previously unstudied proteins capable of programmable transcriptional repression of endogenous genes in mammalian cells.

### CR activity is generalizable across different genes and human cell lines

Next, we sought to determine the broad utility of our newly identified dCas9-CR repressors across different endogenous genes in human cells. First, we selected a subset of CRs that yielded a range of repressive activity from our initial screen. Then, we performed multiplexed repression assays by transiently expressing each dCas9-CR in HEK293T cells that stably express sgRNAs targeting the nonessential genes *CD55* and *CD81* (Figure S2A). We simultaneously quantified CD55 and CD81 protein levels by antibody staining five days post-transfection of the dCas9-CRs (**Figure 4H** and S2B-C). SAP25_HUMAN, MBD3_HUMAN, and RCOR1_HUMAN consistently induced simultaneous strong repression of CD55 and CD81. Other CRs such as WDR82_HUMAN and DPY30_HUMAN induced more intermediate levels of silencing relative to dCas9 alone and CRISPRi. Although human CRs generally outperformed non-human repressors, some non-human CRs such as CMT3_ARATH, A0A921RMW1_SORBI and CHZ1_YEAST also induced repression albeit at weaker levels.

To assess repression strengths across CRs and gene targets in human cells, we normalized fold-change measurements per gene between 0 to 1 to enable comparison of relative repression strength across CRs and visualized these values as heatmaps (**Figure 4I**). This analysis highlighted broadly effective repressors such as RCOR1_HUMAN and SAP25_HUMAN that consistently produced strong repression, along with more gene specific CRs like MTA2_HUMAN and A0A921RMW1_SORBI. These results show that repressors identified in the initial arrayed screen can silence different endogenous gene targets, highlighting their general repressive activity in human cells.

We next sought to evaluate the generalizability of our newly identified CR repressors across human cell types. First, we selected CRs that induced repression in HEK293T cells, including SWC2_ARATH, SAP25_HUMAN, WDR82_HUMAN, RCOR1_HUMAN, and we assessed their activity against the endogenous *CD81* gene in K-562 leukemia cells. We generated stable cell lines each co-expressing a dCas9-CR fusion and sgRNAs targeting the *CD81* promoter or non-targeting sgRNA negative controls. We quantified CD81 protein levels by single cell flow cytometry (**Figure 5A**). Our results show conservation of repressive activity of CRs across cell types as we measured robust repressive activity for SAP25_HUMAN, RCOR1_HUMAN, and DPY30_HUMAN (**Figure 5B**). SAP25_HUMAN and RCOR1_HUMAN showed comparable repression strength to CRISPRi, while DPY30_HUMAN repression was intermediate to dCas9 and CRISPRi. Meanwhile, CRs from non-human organisms basally silenced gene expression similar to dCas9 alone. Taken together, these findings indicate that the repressive activity of CRs is broadly retained across different human cell types.

**Figure 5.**
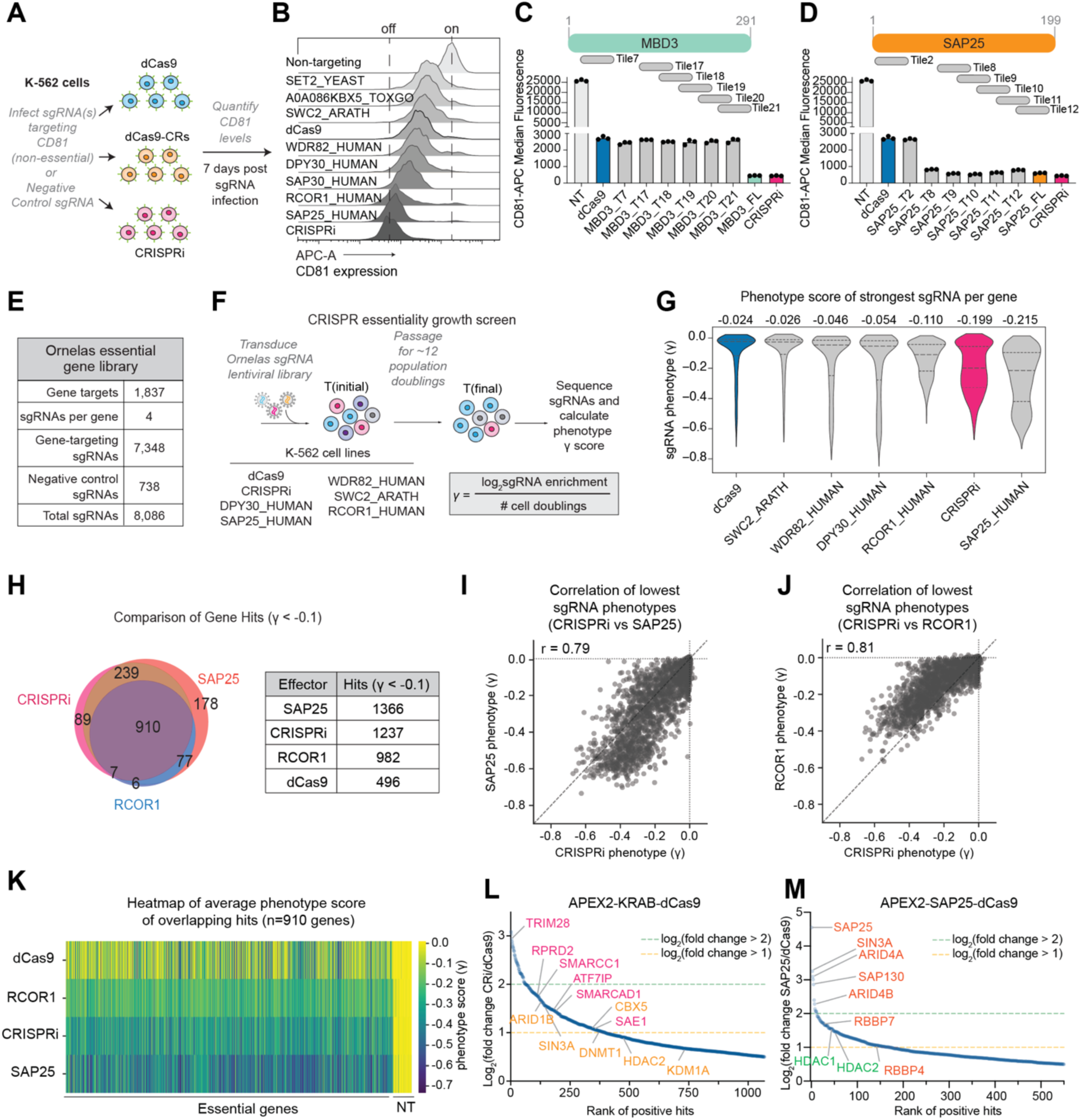
Scalable tuning of cellular phenotypes with CR repressors. **(A)** Schematic of targeting endogenous gene, CD81, in K-562 cells stably expressing sgRNAs with antibody staining for dCas9-CR fusions. **(B)** Flow cytometry histogram plots showing CD81 antibody staining in cells expressing selected CRs 7 days post infection. **(C)** Median fluorescence intensities of CD81 repression for full-length MBD3 compared to corresponding domains. **(D)** Median fluorescence intensities of CD81 repression for full-length SAP25 compared to corresponding domains. **(E)** Table detailing the sgRNA library composition of the Ornelas essential gene library. **(F)** Schematic of pooled CRISPR-based growth screening strategy in K-562 stably expressing dCas9-CR fusions. **(G)** Violin plots of the average lowest gene phenotype scores across CRs from two biological replicates. **(H)** Venn diagram comparing the number of gene hits between select CRs from the essential gene screen. **(I)** Scatter plot showing correlation of average phenotype scores between SAP25_HUMAN and CRISPRi. **(J)** Scatter plot showing correlation of average phenotype scores between RCOR1_HUMAN and CRISPRi. **(K)** Heatmap of average phenotype scores (γ) for overlapping essential gene hits (n = 910). Each column represents a gene and each row an effector; lower scores reflect stronger repression. **(L)** Ranked enrichment plots of fold change of identified proteins from APEX2-KRAB-dCas9 (CRISPRi) compared to APEX2-dCas9 alone. The annotated protein hits are interactors of KRAB and TRIM28 (pink) and known heterochromatin-associated factors. **(M)** Same as in **(L)** but for APEX2-SAP25-dCas9 compared to APEX2-dCas9. The annotated hits are known interactors of the SIN3A-HDAC complex.

Previously, large-scale efforts to test CR functionality relied on 80 amino acid fragments instead of full-length proteins. To compare the repressor efficacy of full-length proteins with fragments, we compared the repressive activity of full-length CRs to their respective ∼80 amino acid tiles previously identified as repressors in published screens^20^. We generated six 80 amino acid fragments for MBD3 and SAP25 – two of the strongest repressors from our screen – and tested their activity on *CD81* in K-562 cells that are stably expressing dCas9 fusions. Using CD81 antibody staining, we observed that full-length MBD3 exhibited stronger silencing than its corresponding tiles, whereas four of the five SAP25 tiles performed similarly to the full-length protein (**Figures 5C-D**). These findings emphasize the advantage of using full-length CRs to capture their native biochemical properties and potential endogenous protein interaction partners, features that may be missed in tile-based approaches.

### A toolkit for programmable titration of cellular phenotypes in human cells

Our identification of repressors that tune gene expression at distinct levels yields potentially new platforms for scalable titration of gene expression and cellular phenotypes in human cells. To test the generality of our newly identified CRs, we performed pooled CRISPR screens to target genes that are essential for cell proliferation in K-562 cells. Though cellular growth is non-linear, it can be used as a proxy for comparing repression strengths of CRISPR perturbation platforms^79^.

We compiled a set of ∼1,800 genes classified as essential for cancer cell viability based on the Cancer Dependency Map (DepMap)^80^. Since our initial arrayed screens assessed CR activity using one sgRNA per target gene, we profiled the broader repressive range of each CR by selecting four sgRNAs from previously published CRISPRi libraries. We included sgRNAs targeting non-essential genes and non-targeting controls, resulting in a final library of 8,086 sgRNAs that we term the Ornelas essential gene library (**Figures 5E-F** and Table S5). We transduced the Ornelas library into K-562 cells expressing dCas9, CRISPRi, SAP25_HUMAN, RCOR1_HUMAN, DPY30_HUMAN, WDR82_HUMAN, or SWC2_ARATH, passaged the cells for ∼12 population doublings, and calculated a γ score by comparing sgRNA abundance at the final time point (Tfinal) to the initial population (Tinitial).

We first compared the distribution of γ scores across the seven CRISPR editors. Across all editors, we observed strong dropout for sgRNAs targeting essential genes, while non-essential and non-targeting controls remained largely unchanged, confirming that our assay effectively depletes essential genes over the time course (Figures S3 and S4A). Notably, SAP25 emerged as a potent and broad repressor, producing strong growth phenotypes across the sgRNA library. Volcano plots comparing gene-level phenotype scores (average of top three sgRNAs per gene) and associated Mann–Whitney p-values yielded thousands of statistically significant genes below a threshold for SAP25 (n= 1,095 genes) and CRISPRi (n= 1,229 genes), highlighting their robust repressive activity across over a thousand essential genes (Figures S4B-D).

To directly compare the targeting profile across editors, we applied an empirically defined cutoff (γ < –0.1) to assess repression of essential genes using the strongest sgRNA per gene for each editor. Violin plots of the lowest phenotype score per gene showed that SAP25 silenced a greater number of genes than CRISPRi at this threshold, outperforming other candidates including RCOR1, WDR82, and DPY30 (**Figure 5G**). Interestingly, although DPY30 and WDR82 are canonical activators, they nonetheless produced widespread depletion of essential gene guides. The overlap analysis of gene hits revealed that SAP25 and CRISPRi shared the majority of hits (∼79% gene hit overlap), but both also have unique targets that highlights their complementary use in functional genomics applications (**Figure 5H**).

A comparison of per-gene phenotypes between SAP25 and CRISPRi showed a strong correlation (r = 0.79) but with a noticeable downward shift towards SAP25, indicating SAP25 as a more potent repressor than KRAB-based CRISPRi (**Figure 5I**). Similarly, comparison of RCOR1 to CRISPRi revealed a strong correlation (r = 0.81) but with RCOR1 consistently producing intermediate phenotypes (**Figure 5J**). We next examined the 910 overlapping essential gene hits between RCOR1, SAP25, CRISPRi, and dCas9 alone, which yielded a gradient of phenotype scores across the CRs. Specifically, the median phenotype score for dCas9 was –0.037, whereas RCOR1, CRISPRi, and SAP25 induced stronger phenotypes with scores of –0.1467, –0.2513, and –0.2859, respectively (**Figure 5K**). Together, these results show that different CRs generate distinct levels of growth defects across a large set of essential genes.

Importantly, among sgRNAs targeting the same gene, different CRs produced different levels of repression for the same sgRNA. This finding suggests that the dynamic repression strengths we observe are due to the CRs themselves and not an artifact of sgRNA choice, highlighting the robustness of the tunable phenotypes with these CRs (Figures S4E-F). Collectively, our pooled screening results establish SAP25 as a robust programmable repressor in human cells, capable of silencing thousands of endogenous genes with strength comparable to CRISPRi. Additionally, using cellular growth as a measurable cellular phenotype, these analyses demonstrate our development of a CR toolkit that can fine-tune phenotypes at defined levels in human cells.

### Proteomic characterization of SAP25 interaction partners

Our identification of SAP25_HUMAN as a strong repressor motivated us to dissect its mode of repression by identifying its protein interaction partners in cells. We performed proximity-based labeling by fusing APEX2 to dCas9, KRAB and SAP25, enabling biotinylation of nearby proteins after effector recruitment to our reporter gene *CLTA-GFP* (Figure S5A). Cells were transfected with plasmids encoding APEX2-dCas9, APEX2-KRAB-dCas9 or APEX2-SAP25-dCas9. At 24 hours post-transfection, cells were treated with biotin and biotinylated proteins were purified with streptavidin followed by mass spectrometry analysis to identify interaction partners that mediate the silencing of CLTA-GFP.

We compared the enriched protein factors in CRISPRi and SAP25 to dCas9 alone to identify protein interactions that are specific to KRAB and SAP25 (**Figures 5L-M** and S5B-C). For SAP25, we detected strong enrichment of SIN3A, RBBP4/7, and SAP30 – known components of the SIN3A co-repressor complex – along with HDAC1 and HDAC2, key histone deacetylases that mediate transcriptional repression^71^. APEX2 profiling of CRISPRi recovered the canonical binding partner of KRAB, TRIM28, and heterochromatin proteins, consistent with the recruitment of KRAB-associated factors for transposon silencing in mammals^66^. The distinct set of interactors highlight that SAP25-mediated gene repression is mediated through the SIN3A-HDAC pathway, while CRISPRi recruits KRAB specific proteins, confirming that these CRs function through distinct pathways.

Altogether, our results highlight the utility in our expanded screen for CRs that can program transcriptional tuning of endogenous genes and is transferable across different cell types. By systematically testing full-length CRs from multiple origins, we discover transcriptional modulators such as SAP25 that recruit distinct chromatin-remodeling machinery. Sampling this broader set of CRs uncovered factors with distinct repression profiles that are different from KRAB and more broadly, expands the available toolkit for programmable repression in mammalian cells.

### Identification of universal CRs in the divergent eukaryote *Saccharomyces cerevisiae*

Our CR screens yielded repressors functional in two distinct eukaryotic kingdoms – Plantae and Animalia – suggesting that many CRs can retain functionality when transferred across divergent eukaryotes. We next investigated whether our newly identified CRs could be used as universal repressors, tools that can maintain consistent function and are utilizable across diverse eukaryotic hosts. To identify universal repressors, we further tested select CRs in the model budding yeast *Saccharomyces cerevisiae* (*S. cerevisiae*) which is a representative organism in the Fungi eukaryotic kingdom.

We first filtered for CRs that functioned as repressors in both *N. benthamiana* and HEK293T cells. We identified 77 CRs that functioned as repressors in *N. benthamiana*, 80 repressors in HEK293T cells, and 29 CRs that acted as repressors in plants and human cells (**Figure 6A**). We refer to these 29 shared repressors as our candidate universal repressors. Our candidate universal repressors have diverse functions in chromatin regulation, with histone deacetylation as the most represented (**Figure 6B**). The annotated Pfam domains of the candidate universal repressors relatively matched annotated functions (**Figure 6C**). Notable candidate universal proteins included RCOR1_HUMAN, SAP25_HUMAN, SWC2_ARATH, and the HBO1 complex proteins KAT7_HUMAN, ING4_HUMAN, and JADE1_HUMAN.

**Figure 6.**
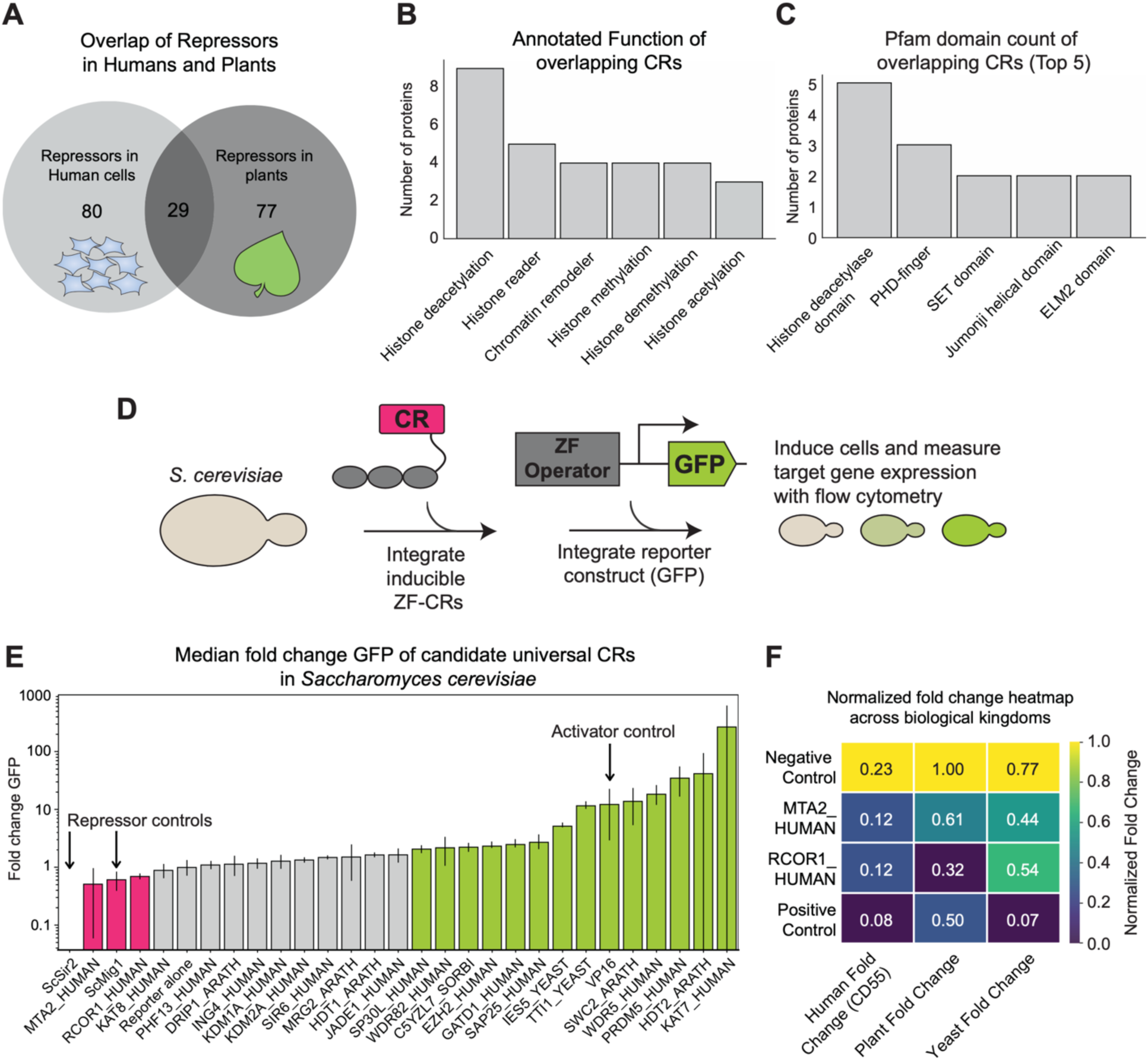
Validation of candidate universal CRs in the divergent eukaryote Saccharomyces cerevisiae. (**A**) Venn diagram showing the overlap of repressors identified in *N. benthamiana* and *H. sapiens*. (**B**) Bar plot showing the annotated function of candidate universal CRs identified as repressors from plant and human cell screen. (**C**) Bar plots showing the top 5 most frequent domains identified for the universal CRs. (**D**) Schematic of the assay workflow for testing transcriptional gene modulation by candidate universal CRs in yeast. (**E**) Bar plot showing the mean fold-change in GFP expression of 3 replicates for CRs tested in the yeast reporter assay. Individual CRs are displayed as bars, with color indicating relative activity (pink, repressors; gray, neutral; green, activators). The reporter alone is the neutral control; ScSir2 and ScMig1 are repressor controls and VP16 is an activator control. Error bars represent the standard deviation. (**F**) Heatmap showing the normalized fold change for universal repressor CRs, MTA2_HUMAN and RCOR1_HUMAN, across biological organisms.

We evaluated the activity for 25 of the 29 candidate universal repressors in *S. cerevisiae* using a previously described synthetic zinc-finger (ZF)-CR screening system^17^. In this system, the ZF-CR is recruited upstream of a minimal CYC1 promoter to modulate GFP expression after chemical induction. In the background strain YPH500, we integrated three cassettes: 1) a constitutive repressor construct that prevents expression of the ZF-CRs until release via the addition of chemical inducers, 2) the fluorescent reporter, and 3) the ZF-CR. To benchmark activity, we used *S. cerevisiae* Mig1 and Sir2 as control repressors and VP16 as an activator control. Correcting for background fluorescence, we defined repressors as CRs producing <0.7 GFP fold change normalized to the reporter alone, and activators as CRs exceeding a fold change of 2.0 (**Figure 6D**).

Given that humans and yeast are more closely related evolutionarily (both within the Opisthokonta eukaryotic supergroup) than humans and plants (Archaeplastida eukaryotic supergroup)^81^, we hypothesized that the candidate universal repressors would retain function in yeast. However, only 2 of the 25-candidate universal CRs – RCOR1_HUMAN and MTA2_HUMAN – induced repressive activity in *S. cerevisiae*, with RCOR1_HUMAN being amongst the strongest repressors in human and plant systems. In contrast, thirteen of the CRs functioned as activators in yeast with five, SWC2_ARATH, WDR5_HUMAN, PRDM5_HUMAN, HDT2_ARATH, and KAT7_HUMAN, showing activation stronger than our activator control VP16, while the remaining CRs were largely neutral. Notably, IES5_YEAST and TTI1_YEAST were previously identified as activators in yeast^17^, highlighting agreement with previous reports and differences from activities in other systems.

Together, these findings show that most CRs have species-specific activity, but a small subset, including RCOR1_HUMAN and MTA2_HUMAN, maintain their repressive activity across evolutionary distant hosts (**Figure 6E**). Their consistent activity across human, plant, and yeast systems suggests a conserved repressive mechanism that can be harnessed for programmable gene modulation across eukaryotes from different biological kingdoms.

## DISCUSSION

Chromatin regulators are highly prevalent across the eukaryotic biological kingdoms with an estimate of >40,000 CRs that exist in available eukaryotic genomes^9^. However, the ability to test their functions in living cells has been limited. Here, we present the largest effort to date experimentally testing the functions of hundreds of full-length CRs in divergent eukaryotic hosts, thereby enabling us to decipher the direct effect of eukaryotic CRs on gene expression.

Our study yielded new technologies for programmable transcriptional modulation in plants and human cells, including new CRs that outperform current state-of-the-art platforms. In plants, our discovery of potent CRs opens opportunities for applications in synthetic biology across a wide range of crops by expanding the available toolkit of trans-elements for genetic engineering. In human cells, we discover CRs that enable tuning of gene expression to defined intermediate levels, including SAP25, a potent repressor that outperforms CRISPRi-KRAB. We further apply our programmable CRs for large-scale tuning of over a thousand genes, thereby allowing us to titrate cell proliferation phenotypes at distinct rates.

Our study highlights how the core repressive activities of CRs can be conserved when heterologously transferred across distinct eukaryotic kingdoms. In particular, repression through histone deacetylation appears to represent a conserved mechanism across eukaryotes, with HDAC-containing complexes being a unifying mechanism for gene silencing in plants, animals and fungi^82^. As such, SAP25 emerged as a potent repressor in human cells, outperforming KOX1-KRAB for functional genomics screens in K-562 cells. SAP25 and other members of the SIN3A complex also functioned as repressors in plants, emphasizing the evolutionary conservation of HDAC-mediated repression across kingdoms. In contrast, the state-of-the-art KRAB repressor in human cells, which harnesses the natural pathway of epigenetic repression of transposable elements in mammals through H3K9me3 deposition, is not a strong repressor in plants (Table S2). More generally, the relatively small overlap of common CRs between plants, humans and yeast sheds light into the context-dependent functions of chromatin regulators across the eukaryotic kingdoms.

Unexpectedly, we observed numerous non-native CRs that retain their functionality when expressed outside of their native host, as seen with C5YUK9_SORBI and P53_HUMAN that produced the strongest activation in humans and plants, respectively. We hypothesize that these CRs may have less regulation when expressed heterologously, thereby enabling them to induce robust transcriptional activation^27^. Our unexpected discovery that C5YUK9_SORBI and P53_HUMAN outperform state-of-the-art synthetic activators in humans and plants highlights the novelty of our approach to experimentally test CRs outside their endogenous context and presents a prime opportunity for synthetic biology, where non-native regulators may offer modular control of gene expression.

Finally, we identified the human-encoded RCOR1 and MTA2 proteins as universal CRs that retain their transcriptional repression activity in plants, human cells, and yeast, thereby signifying their utility for synthetic gene control in divergent eukaryotic hosts. Both are members of established repressive chromatin complexes, the CoREST complex and NuRD complex, respectively, which recruit histone deacetylases^49,83^. Their ability to function as repressors in three distinct eukaryotic kingdoms suggests that parts of these complexes, or their domains, are functionally conserved and can work independent of species-specific cofactors. Although yeast lack canonical NuRD and CoREST complexes, they retain CRs with analogous functions (i.e. deacetylases and remodeling complexes) that may enable these human repressors to function outside of their endogenous host. Our identification of RCOR1_HUMAN and MTA2_HUMAN as universal repressors establishes a set of effectors that offers a toolkit to implement for transcriptional and epigenetic manipulation in non-model organisms where such tools are lacking. These findings highlight how leveraging evolutionary diversity can uncover CRs with conserved activity capable of functioning beyond their native context, enabling gene expression control in diverse hosts for use in biotechnology, agriculture, and therapeutic settings.

## MATERIALS AND METHODS

### Constructing the CR library

To select plant genes for the CR library, we first sought to gather a representative list of *Arabidopsis thaliana* proteins from all major classes of histone modifiers, chromatin remodelers, and proteins involved in DNA methylation, as well as diverse set of proteins involved in epigenetic regulation. For three major classes of histone modifiers (deacetylases, acetyltransferases, and lysine methyltransferases), we selected all the annotated members in the *A. thaliana* TAIR10 genome. For the rest of candidate Arabidopsis CRs, we chose only one candidate when multiple close orthologs were found (>90% sequence identity). These *Arabidopsis* genes included representative histone lysine demethylases of the JMJ class and enzymes involved in other histone posttranslational modifications such as phosphorylation. Beyond histone modifiers, we selected all the *Arabidopsis* members of chromatin remodeler complexes including the CAF1 and FIE complex, Polycomb group (PcG) proteins, ISWI complex and ISWI-associated proteins, the INO80 complex, the SWR1/SRCAP complex, SWI/SNF components, and the NuA4 histone acetyltransferase complex. We also selected known H3K4me3 binding proteins, PHD chromatin readers, histone chaperones, and components of the nuclear pore and nuclear lamina. We also included all the genes known to be involved in the DNA methylation pathway at the level of establishment and readout of this mark. We next used this list of *Arabidopsis* genes to find their *Sorghum bicolor* homologs using sequence similarity. To this end, we used the Phytozome database^84^ and identified the closest *S. bicolor* homolog, regardless of which Sorghum variety the genes belonged to. To select *T. gondii* genes, we followed a similar approach by using representative genes from each of the aforementioned classes to guide a homology-based search using NCBI BLAST. However, unlike for the *S. bicolor* search, we prioritized the use of *H. sapiens* representative proteins for this selection, relying on annotated human CRs using the EpiFactors database^85^. *S. cerevisiae* proteins were selected from previously published datasets that were identified as repressors or activators^17^. Finally, for the selection of *Megaviridae* candidates we picked the longest protein from each major class of CRs found in this taxon. Sequences were codon-optimized for *N. benthamiana* flanked by BsaI sites for downstream cloning (see: cloning the CR library for expression in *Nicotiana benthamiana*).

### Cloning the CR library for expression in *Nicotiana benthamiana*

The PMS7997 backbone {pMAS:GAL4 DBD-GGSGG-GFP dropout:tNOS} was digested with BsaI-HF^®^v2 to excise the GFP dropout. The digested backbone was purified from the gel using Zymogen™ Gel Cleanup Kit. CRs were digested using New England Biolabs^®^ (NEB) BsaI-HF^®^v2 and ligated into the BsaI-HF^®^v2-digested PMS7997 backbone using Thermo Scientific™ T4 DNA ligase. Ligation products were transformed into XL-1 blue *Escherichia coli* competent cells via heat shock, recovered in 1 mL liquid BD Difco™ LB (Luria-Bertani) Broth Miller at 37°C for 1 h, and streaked onto solid LB containing 50 mg/L Kanamycin. Colonies that survived selection were cultured, miniprepped using the QIAGEN plasmid prep kit, and sequenced (primer: 5’-CCTCTAATAAGGGTCAAAGAC-3’). Correct plasmids were electroporated into GV3101 *Agrobacterium tumefaciens* competent cells using the Bio-Rad GenePulser Xcell™ at 2400 V, 25 μF, 200 Ω for 4-5 ms in 1 mm cuvettes. Transformed cells were recovered in 1 mL liquid LB at 30°C for 3h then plated on solid LB with 50mg/L Kanamycin, 30mg/L Gentamicin, and 50mg/L Rifampicin (LB RGK). All plasmids and strains are available from the Joint BioEnergy Institute (JBEI) Inventory of Composable Elements (ICE) registry: (https://public-registry.jbei.org/folders/941; Table S10).

### Nicotiana benthamiana growth conditions

The *N. benthamiana* (accession LAB) plants were grown in a room at 25°C with 16h light: 8h dark cycle at 150 μmol m^−2^ s^−1^ photosynthetically active radiation (PAR; wavelength: 400–700 nm) and 43% humidity. Seeds were planted in water-saturated Sunshine MIX #4 soil supplemented with 1 Tablespoon/2 gallons Osmocote Classic 14-14-14 covered with ∼3 cm CR Premier Pro-Mix PGX topsoil in pots placed in a tray with a vented hood with the vents closed. After one week, seedlings were transplanted to individual pots without topsoil with eighteen pots in each tray. After the soil was saturated with water, excess water was drained, and the seedlings were transplanted. Vented hoods with the vents closed were placed on each tray. At two weeks of age, the vents of the hood were opened. At three weeks of age, the hoods were removed, and each tray received 1 L water supplemented with 1 teaspoon/gallon Peter’s Professional 20-20-20. At three weeks and five days, the plants were split into nine pots per tray and each tray received 1.5 L water supplemented with 1 teaspoon/gallon Peter’s Professional 20-20-20. At 4 weeks of age, the *N. benthamiana* plants were agroinfiltrated.

### Nicotiana benthamiana agroinfiltration

Transient expression is accomplished via “agroinfiltration”––pushing *Agrobacterium tumefaciens* cells into the apoplast of plant leaves where they can transfer DNA into the plant cells^86^. The day before agroinfiltration, excess water was drained from the plant trays and *A. tumefaciens* GV3101 strains containing either the reporter plasmid, PMS6370 {5X GAL4 binding sites-pcoreWUSCHEL:GFP:tNOS}, or a CR plasmid {pMAS:GAL4 DBD-GGSGG-CR:tNOS} were cultured in liquid LB RGK. Cultures were grown at 30°C, shaking at 200 rpm for 18 h. After 18 h, the cultures were diluted in 4 volumes of LB RGK and grown at 30°C for 1-2 h or until the optical density at 600 nm (OD_600_) was between 0.5-2.0. The cultures were spun down at 3220 x g for 15 min. The LB was removed, and the agrobacteria resuspended with 1X infiltration buffer (10 mM magnesium chloride (MgCl_2_), 10 mM 2-(N-morpholino)ethanesulfonic acid (MES), 0.2 mM acetosyringone, pH 5.6). The volume of infiltration buffer to resuspend the agrobacteria in was calculated using the following formula:

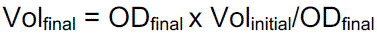

The agrobacteria were placed on a rocking shaker for 1.5 h before infiltration. The reporter and CR strains were diluted to a final OD of 0.5. Individual CR strains were mixed with the reporter strain at a 1:1 ratio. Using a blunt syringe, each strain mix was infiltrated into two leaves of a plant for three plants per mix. The first two fully expanded leaves were infiltrated, approximately the fourth and fifth leaves counting from the shoot apical meristem (SAM). After agroinfiltration, the plants were placed back onto their tray and put back into the growth room. The day after agroinfiltration, each tray of agroinfiltrated plants received 1 L water.

### *N. benthamiana* fluorescence data collection

Leaf fluorescence data was collected at 4 days post-infiltration (dpi). Using a hole punch, four samples of leaf tissue (“leaf punches”) were collected from each infiltrated leaf, typically two from each side of the leaf. Veins and syringe marks were avoided. Leaf 1 was considered the younger leaf (closer to the SAM) and leaf 2 was the older leaf (one node further from the SAM). Plants were labeled 1, 2, or 3 and any abnormalities were recorded, such as chlorosis or necrosis around the infiltration zone. Each well of a 96-well plate with a clear bottom was filled with 350 μL tap water and, using forceps, each leaf punch was placed on top of the water in a well with the abaxial side of the leaf facing up. Using a BioTek Synergy H1 Microplate Reader, the fluorescence of each leaf punch was measured for GFP (Ex: 488 nm, Em: 520 nm).

### *Nicotiana benthamiana* agroinfiltration data analysis

To calculate fold change GFP, the mean GFP fluorescence of each CR was divided by the mean GFP fluorescence of the reporter strain co-infiltrated with a plasmid containing the Gal4 DBD but no CR (GAL4 DBD alone). As each week was functionally an independent experiment, all controls (GAL4 DBD alone, VP16, and SUPERMAN) were repeated each week, and each week’s data was normalized to the Gal4 DBD infiltration performed that week.

### Human cell culture

All cell lines were obtained and authenticated by the UC Berkeley Cell Culture Facility. The CLTA-GFP and RAB11A-GFP HEK293T cell lines originated from a previous study^58,59^. HEK293T cells were cultured in DMEM (Gibco). K-562 cells were cultured in RPMI-1640 (Gibco). All media was supplemented with 10% (v/v) FBS (VWR) and 100 U/mL streptomycin, 100mg/mL penicillin (Gibco). Cell lines were cultured at 37°C with 5% CO_2_ in tissue culture incubators and routinely confirmed to be negative for *Mycoplasma*.

### Cloning the CR library for expression in human cells

The pIZ48 backbone {pCAG:dCas9-TagBFP-GFP dropout} was digested with NEB PaqCI to excise the GFP dropout. The digested backbone was purified from the gel using Zymogen™ Gel Cleanup Kit. CRs were digested using NEB BsaI-HF^®^v2 and ligated into the PaqCI-digested pIZ48 backbone using NEB T7 DNA ligase. Ligation products were transformed into Stellar *Escherichia coli* competent cells (Takara Bio) via heat shock, recovered in 0.2 mL liquid (Luria-Bertani) Broth Miller at 37°C for 1 h and streaked onto solid LB containing 100 mg/L Carbenicillin. Colonies that survived selection were cultured, miniprepped using the Qiagen plasmid prep kit, and sequenced (primer: 5’-AGGAGGCCAACAACGAGACCTACG-3’).

### Generation of stable sgRNA expressing cell line and lentiviral production

To test the dCas9-CRs in human cells, HEK293T and K-562 cell lines were engineered to stably express sgRNAs that target various endogenous or reporter loci. In HEK293T cells, sgRNAs targeting the *CLTA* and *RAB11A* promoters were introduced into CLTA-GFP and RAB11A-GFP reporter lines, respectively. Additionally, sgRNAs targeting endogenous genes *CD55* and *CD81* were delivered to generate stable knockdown lines. In K-562 cells, sgRNAs targeting the *CD81* promoter were stably expressed.

Lentiviral particles were produced by transfecting standard packaging vectors into HEK293T using TransIT®-LT1 Transfection Reagent (Mirus Bio, MIR2306). Cells were seeded at a density of 4.0 × 10^5^ cells per well in a 6-well BioLite Microwell plate (Thermo Fisher Scientific) and co-transfected the next day with standard packaging plasmids (0.1 μg each of gag-pol, REV, and TAT; 0.2 μg VSVG) and 1.5 μg of the lentiviral sgRNA expression vector (Addgene #217306) using Opti-MEM (Gibco) and TransIT®-LT1 Transfection Reagent (Mirus Bio), following the manufacturer’s instructions. Media was changed 24 hours post-transfection with complete DMEM. Viral supernatants were collected 48–60 hours post-transfection and filtered through a 0.45 μm PES syringe filter (VWR). Target cells were transduced with lentiviral particles and sorted based on mCherry expression to enrich for sgRNA-expressing populations, resulting in cell lines with >90% sgRNA expression (mCherry positive).

### dCas9-CR transfections and analysis

Transient transfection experiments in HEK293T cells were performed in 96-well plates using TransIT®-LT1 (Mirus Bio) and Opti-MEM™ I Reduced Serum Medium (Thermo Fisher Scientific). Cells at ∼70–80% confluency were transfected with 150 ng of plasmid encoding each dCas9-CR construct. Transfection efficiency was measured by BFP fluorescence, with a minimum cutoff of 20% BFP+ cells required for inclusion in downstream analysis. Cells were collected at multiple time points: Day 2, 5, 7, 10, and 14 post-transfection for flow cytometry analysis. Relative activity was calculated as: (% GFP-cells on Day 2 / transfection efficiency) × 100. To evaluate the strength of silencing, fold change was computed as: (Day 5 median GFP intensity in the GFP-gate / Day 5 median GFP intensity in untransfected cells).

To evaluate the dCas9-CRs for multiplexed repression of CD81 and CD55, we first engineered HEK293T cell lines to stably express sgRNAs targeting *CD55* and *CD81.* Cells were seeded in 24-well plates and transfected at ∼70–80% confluency with 300 ng of plasmid encoding select dCas9-CRs using TransIT®-LT1 Transfection Reagent and Opti-MEM™ I Reduced Serum Medium, according to the manufacturer’s protocol. At 5 days post-transfection, cells were harvested and stained with fluorophore-conjugated antibodies specific to CD55 and CD81. Surface protein expression in single cells was assessed by flow cytometry. Gates were defined based on isotype and untransfected controls to quantify silencing relative to baseline expression.

Quantification of CD2, CD55 and CD81 protein levels were measured by cell surface antibody staining of live cells. Cells were incubated with APC– or FITC-labeled antibody (CD2: Miltenyi Biotec/ CD55 and CD81: BioLegend) for ∼30 min in the dark at room temperature, washed twice with PBS containing 10% FBS, and protein expression was measured on a BD FACSymphony A1 Cell Analyzer (BD Biosciences) flow cytometer. All flow cytometry data generated was analyzed using FlowJo.

### Essential gene screen

For the growth-based screen targeting essential genes in K-562 cells, an sgRNA library was designed based on a previously published genome-wide CRISPRi screen in K-562 cells and the DepMap CRISPR-inferred common essential ^80,87^. The library consists of 8,086 sgRNAs targeting 1,838 essential genes, along with non-essential and non-targeting control sgRNAs. Each essential gene is targeted by four sgRNAs, selected from previously published sgRNA library^64,87^. A list of protospacer sequences included in the essential-gene library is available in Table S5.

sgRNA sequences were cloned into an expression plasmid obtained from Addgene (ID#217306), which includes a U6 Pol III promoter and a T2A-mCherry reporter to measure transduction efficiency. The sgRNA oligos were synthesized as a pooled library (Twist Bioscience) with the structure: 5’ – PCR adaptor – CCACCTTGTTGG – protospacer sequence – GCTAAGC – PCR adaptor – 3’. Oligo pools were PCR-amplified, digested with BstXI/BlpI, gel extracted, and ligated into the sgRNA lentiviral vector. The resulting plasmid library was transformed for amplification and validated by next-generation sequencing to confirm library representation and uniformity.

To conduct the screen, K-562 cells stably expressing a dCas9 fusion protein were transduced with the lentiviral sgRNA library by spinfection (1000 × g for 2 hours at 33°C) with polybrene (8 µg/mL, Sigma-Aldrich). Infection efficiency was assessed two days post-infection by flow cytometry, aiming for 20%–30% mCherry-positive (sgRNA-expressing) cells. Screens were performed with two technical replicates, maintaining a minimum representation of at least 1,000 cells per sgRNA throughout the experiment. Two days post-transduction, puromycin was added to the media until ∼90% of cells were sgRNA-positive, as indicated by mCherry fluorescence. After this, cell pellets were collected and frozen for each replicate to represent the initial time point (T_initial_). Cell doublings were tracked throughout the experiment, and final samples (T_final_) were collected at approximately 12 cell doublings.

DNA libraries for T_initial_ and T_final_ were prepared for deep sequencing following a previously described protocol^64^. Genomic DNA was extracted from cell pellets using the NucleoSpin Blood L kit (Macherey–Nagel). The genomic DNA was used as template for directPCR-amplification of the protospacer sequence (23 cycles) using NEBNext Ultra II Q5 PCR Master Mix (NEB) using primers with appended Illumina adapters and unique sample indices. Libraries were sequenced on a NovaSeqX platform with a 19 bp Read 1 and a 5 bp Index 1.

### APEX proteomic labeling

HEK293T cells with stable expression of sgRNAs targeting the endogenous *CLTA-GFP* promoter were cultured in 15 cm dish and transfected with plasmids expressing APEX2-SAP25-dCas9, APEX2-KRAB-dCas9 or APEX2-dCas9 only. 24 hours post-transfection, cells were treated with 500 µM biotin-tyramide (MilliporeSigma – SML2135) for 30 min. 1 mM final H_2_O_2_ was added to each dish for a total labeling time of 1 minute. Cells were quenched with 2X quenching buffer (10 mM Trolox and 20 mM sodium ascorbate in DPBS), collected and washed twice with cold DPBS. Cell pellets were lysed with 500 µL cytoplasmic lysis buffer (10 mM Tris pH 8, 1.5 mM MgCl2, 10 mM KCl, 0.2% NP-40, and 1mM DTT) for 10 min on ice. After centrifugation at 2,500 x g for 5 min at 4°C, the pellets were collected and washed once with cold PBS. Cell pellets were lysed in 200 µL nucleus lysis buffer (20 mM Tris pH 8, 25% glycerol, 400 mM NaCl, 1.5 mM MgCl2, 10 mM EDTA, 1 mM DTT, and 0.2 µL benzonase (Sigma-Aldrich, E1014) and incubated on ice for 30 min. After centrifugation at 20,000 x g for 10 min at 4°C, the supernatant was collected, combined with 800 µL of cold RIPA lysis buffer (110% glycerol, 25 mM Tris-HCl pH 8, 150 mM NaCl, 2 mM EDTA, 1% NP-40, 0.2% sodium deoxycholate, 5 mM Trolox, 10 mM sodium ascorbate, and 1x Protease Inhibitor), and incubated with Streptavidin beads (Pierce) at RT for 1 hr. Beads were washed twice with RIPA lysis buffer, once with 1 M KCl solution, once with 2 M Urea, pH 8, and twice with RIPA lysis buffer (without detergent or quenching reagents). Beads were washed three times with ABC buffer (50 mM ammonium bicarbonate, pH 8), rotated head-over-head at 4°C for 20 min. Beads were resuspended in 25 µL ABC buffer with 2.5 µL of trypsin (1 µg/µL) and incubated for 5 hours at 37°C. Peptides were collected, quantified with Fluorescent Peptide Analysis, and 0.6 µg of each sample was subjected to mass-spectrometry (MS)-based analysis. DIA data was analyzed using Spectronaut 18.4 (Biognosys Schlieren, Switzerland) using the direct DIA workflow with PTM localization selected. We identified 171 hits showing the most significant enrichment in APEX2-SAP25-dCas9 and 400 hits showing the most significant enrichment in APEX2-KRAB-dCas9 (Table S7 and Table S8).

### Cloning the CR library for expression in *Saccharomyces cerevisiae*

Yeast expression plasmids were a gift from the Khalil lab and are as previously described in their paper^17^. We used the background strain YPH500 (*MAT*α*/ura3-52/lys2-801_amber/ade2-101_ochre/trp1-*Δ*63/his3-*Δ*200/leu2-*Δ*1*) and genomically integrated all plasmids. The yeast cells were transformed using the LiAc/SS carrier DNA/PEG method. A 5 mL culture of yeast was grown in YPD (1% yeast extract, 2% peptone, 2% dextrose) at 30°C, shaking at 200 rpm, overnight. The next day, 45 mL of YPD was inoculated with the overnight yeast culture and grown at 30°C, shaking at 200 rpm, until reaching an OD_600_ of 0.6-2.0. The cells were spun down at 3000 x g for 5 min and resuspended in 25 mL sterile dH_2_O. The cells were spun down again at 3000 x g for 5 min, resuspended in 1 mL 100 mM lithium acetate (LiAc), and aliquoted into 1.5 mL microcentrifuge tubes of 100 µL. The cells were incubated at 30°C for 15 minutes, shaking at 200 rpm, then spun down at 3000 x g for 30 s. After discarding the supernatant, 30 µL of linearized plasmid (∼1 – 5 µg DNA) was added to the cells. Next, 335 µL of the freshly made T-mix (36% PEG 3350, 107 mM LiAc, 0.3 mg/mL boiled sheared salmon sperm DNA) was added to the cells and vortexed vigorously. The mix was incubated for 30 min at 30°C, shaking at 200 rpm, before incubation at 42°C in a water bath for 60 min. The mix was spun down at 1000 x g for 2 min to pellet the cells. After removing the supernatant, the cells were resuspended in 200 µL sterile dH_2_O, plated on YPD, and grown at 30°C. After colonies appeared, each plate was replica plated with velvets onto selective media and grown at 30°C until colonies grew.

The constitutive repressor construct was transformed into YPH500 and selected on YPD with 200 mg/L Hygromycin B (Thermo Fisher Scientific). The fluorescent reporter was transformed into the constitutive repressor strain and selected on SD media without uracil (0.65 g/L CSM-His-Leu-Ura (Sunrise Science Products 1015-010), 1.7 g/L Yeast Nitrogen Base Without Amino Acids and Ammonium Sulfate (Millipore Sigma Y1251), 38 mM ammonium sulfate, 1.14 mM L-Leucine, 450 µM L-Histidine (0.1M HCl), 200 mg/mL Hygromycin B, 2% dextrose). The ZF-CRs were transformed into the strain with both the reporter and constitutive repressor and selected on SD media without uracil and histidine (0.65 g/L CSM-His-Leu-Ura, 1.7 g/L YNB Without Amino Acids and Ammonium Sulfate, 38 mM ammonium sulfate, 1.14 mM L-Leucine, 200 mg/mL Hygromycin B, 2% dextrose), except for the off-target 97-4 ZF-VP16 with was selected on SD media without uracil and leucine (0.65 g/L CSM-His-Leu-Ura, 1.7 g/L YNB Without Amino Acids and Ammonium Sulfate, 38 mM ammonium sulfate, 450 µM L-Histidine (0.1M HCl), 200 mg/mL Hygromycin B, 2% dextrose).

The constitutive repressor construct (TetR, LacI, and GEV in pNH607) was linearized with PmeI (NEB) and integrated into the HO locus. The constitutive repressor prevented expression of the ZF-CR until release by addition of the chemical inducers anhydrotetracycline (1.0 ug/mL), IPTG (20 mM), and β-estradiol (100 nM). The fluorescent reporter (minimal pCYC1:yEGFP with upstream 43-8 ZF binding sites in pRS406) was linearized with StuI (Eco147I; Thermo Fisher Scientific) and integrated into the URA3 locus. The 43-8 ZF-CRs (ZF amino acid sequence: GAGTGAGGA; in pNH603) were linearized with AflII (Thermo Fisher Scientific) or SalI (Thermo Fisher Scientific) depending upon available cutsites. The ZF-CRs were integrated into the HIS3 locus. Of the 29 candidate universal repressors, five were unable to be cloned: EHMT2_HUMAN, MBD3_HUMAN, PHF8_HUMAN, SUS1_YEAST, and A0A1B6PTU2_SORBI.

### Induction of the CRs in *S. cerevisiae*

After transformation with the above-described cassettes, as available, three individual colonies were used to inoculate 500 µL SD media (synthetic dropout media with 2% glucose and amino acid compositions varying based on necessary selection) in Costar 2 mL 96-well assay blocks. Cultures were grown for 24-48 h at 30°C, shaking at 200 rpm. Next, 5 µL of each culture was used to inoculate 500 µL SD-complete media with or without the previously described chemical inducers and grown at 30°C for 12 h. 10µg/mL cycloheximide was added to inhibit protein synthesis before the cells were assayed via flow cytometry.

### *S. cerevisiae* flow cytometry and data analysis

We set the event collection to 10,000 on a BD FACSymphony™ A1 Cell Analyzer and collected yEGFP fluorescence of the cells (Ex: 485 nm, Em: 510 nm). We gated the events using forward and side scattering and then calculated the median fluorescence of the induced and uninduced cells per strain per replicate. We subtracted the median autofluorescence of the induced and uninduced *S. cerevisiae* YPH500 (no integrations) strain from the median induced and uninduced fluorescence of the transformed strains, respectively. For each strain replicate, we divided median fluorescence of the induced cells by the median fluorescence of the uninduced cells. We then calculated the mean fold change GFP of the three replicate medians to obtain the fold change GFP value.

## ACKNOWLEDGEMENTS

We thank members of the Shih and Nuñez labs for thoughtful advice, discussions, and resources. We are grateful to Ben Williams for the thoughtful discussions. We thank Albert J. Keung and Ahmad ‘Mo’ S. Khalil for the plasmids used in the yeast experiments. We thank Donat Wulf for technical assistance. We thank the UCSF CAT and UCB QB3 genomics (supported by UCSF PBBR, RRP IMIA, and NIH 1S10OD028511-01 grants) for assistance with Illumina sequencing. This work was part of the DOE Joint BioEnergy Institute (https://www.jbei.org) supported by the U. S. Department of Energy, Office of Science, Office of Biological and Environmental Research, through contract DE-AC02-05CH11231 between Lawrence Berkeley National Laboratory and the U.S. Department of Energy. The work (proposal: 505883 DOI:10.46936/10.25585/60000931) conducted by the U.S. Department of Energy Joint Genome Institute (https://ror.org/04xm1d337), a DOE Office of Science User Facility, is supported by the Office of Science of the U.S. Department of Energy operated under Contract No. DE-AC02-05CH11231. J.K.N. acknowledges funding for this work from the Bakar Fellows Program (UCB 51474) and a Hanna Gray Fellowship from the Howard Hughes Medical Institute (GT15341). J.K.N. is funded as an Investigator of Biohub, San Francisco. I.J.O. is funded by a University of California, Berkeley Mentored Research Award, Graduate Research Fellowship from the National Science Foundation, and a Gilliam Fellowship from the Howard Hughes Medical Institute. At the time of the study, L.A.O. was supported by the National Institutes of Health-funded Genetic Dissection of Cells and Organisms Training Program (project number: 2T32GM132022-06) and the Arnon Fellowship from the University of California, Berkeley’s Department of Plant and Microbial Biology. N.P. acknowledges support from a University of California, Berkeley Mentored Research Award and a University of California Chancellor’s Fellowship. R.K.P. is funded by graduate fellowships from the University of California Cancer Research Coordinating Committee and the Shurl and Kay Curci Foundation.

## AUTHOR CONTRIBUTIONS

I.J.O., S.A., P.M.H., N.F.C.H, and J.K.N. curated the chromatin regulator library. I.J.O, L.A.O, P.M.H. and J.K.N. led the study, designed the experiments, and wrote the manuscript with assistance from the co-authors. L.A.O. led the plant and yeast experiments, while I.J.O. performed the majority of the human experiments. M.G.H. assisted with the arrayed screen in human cells and performed the experiments for comparing domains to full-length proteins as transcriptional effectors. N.P. performed the APEX2 proximity labeling experiments and downstream proteomics analysis. R.K.P. assisted with the essential gene screen in K-562s.

## RESOURCE AVALABILITY

All plasmids containing CR sequences can be accessed from the JBEI public repository (https://public-registry.jbei.org/folders/941; Table S10). Plasmids encoding C5YUK9_SORBI, SAP25_HUMAN, RCOR1_HUMAN, and MTA2_HUMAN fused to dCas9 will be deposited on Addgene upon publication. The CR library sequences are available within the supplementary table. All other data are available in the manuscript or the supplementary materials.

---

**Figure S1.**
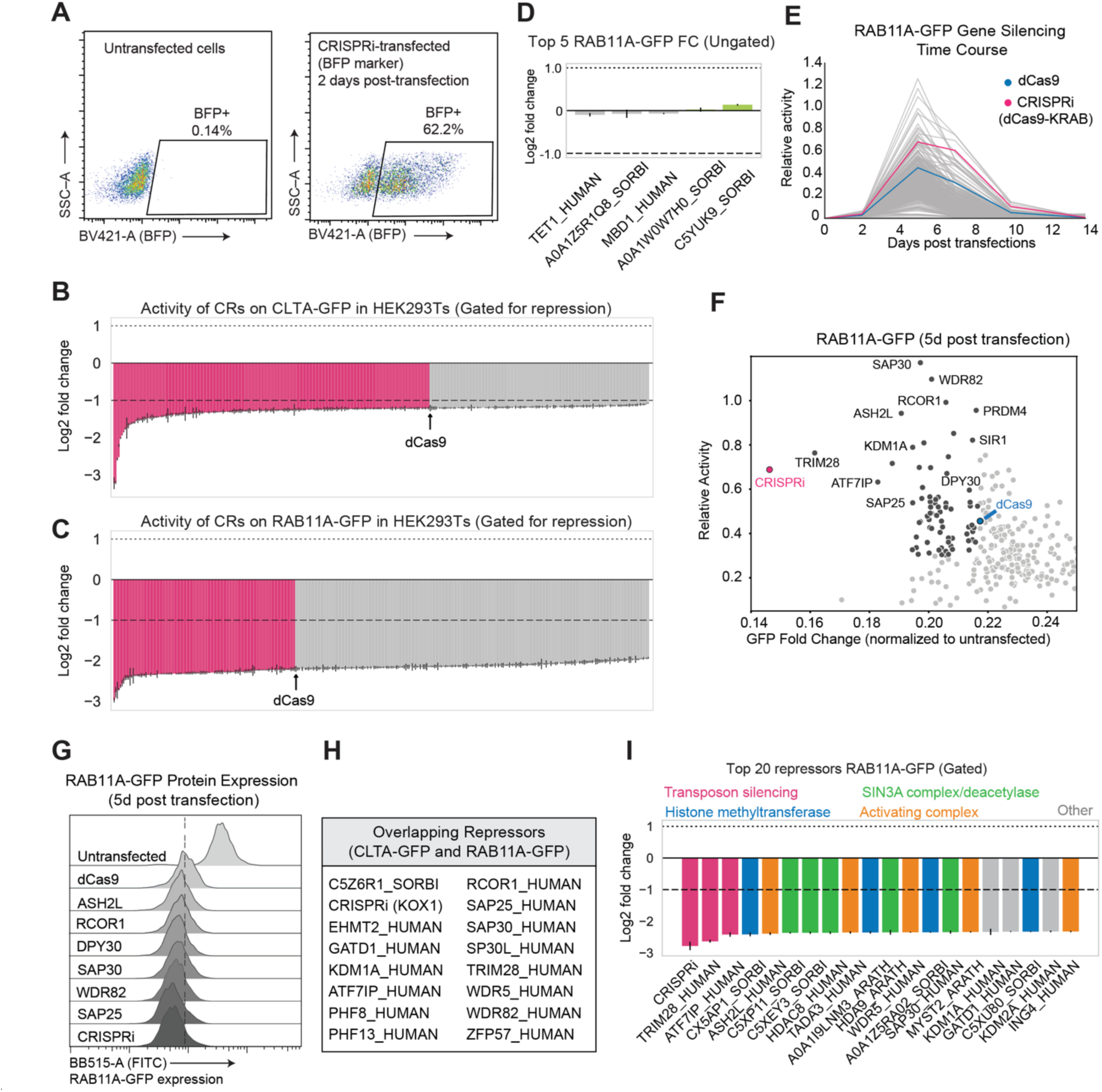
Gated analysis of CR activity in human cells and RAB11A-CR screen activity. **(A)** Representative flow cytometry plots showing transfection efficiency using a BFP marker in untransfected and CRISPRi-transfected HEK293T cells. **(B)** Gated analysis of CLTA-GFP fluorescence to quantify repressive activity of CRs 5 days post-transfection. Each bar represents the log₂ fold-change in reporter expression relative to untransfected controls, with repressors shown in pink. **(C)** Same as in **(B)** but for RAB11A-GFP reporter in human cells 5 days post-transfection. **(D)** Top five-fold-changes for RAB11A-GFP from the ungated analysis. **(E)** Time course of GFP silencing for RAB11A-GFP reporter across tested CRs. **(F)** Scatterplot of relative activity and GFP fold change for RAB11A-GFP at 5 days post transfection. **(G)** Representative GFP distributions from flow cytometry for selected CRs in RAB11A-GFP. **(H)** Uniprot IDs for overlapping repressors identified from CLTA-GFP and RAB11A-GFP. **(I)** Bar plot showing the RAB11A-GFP log₂ fold-change for top 20 repressors identified from human screen for the gated population. Fold changes were calculated using median GFP from GFP-gate compared to untransfected control.

**Figure S2.**
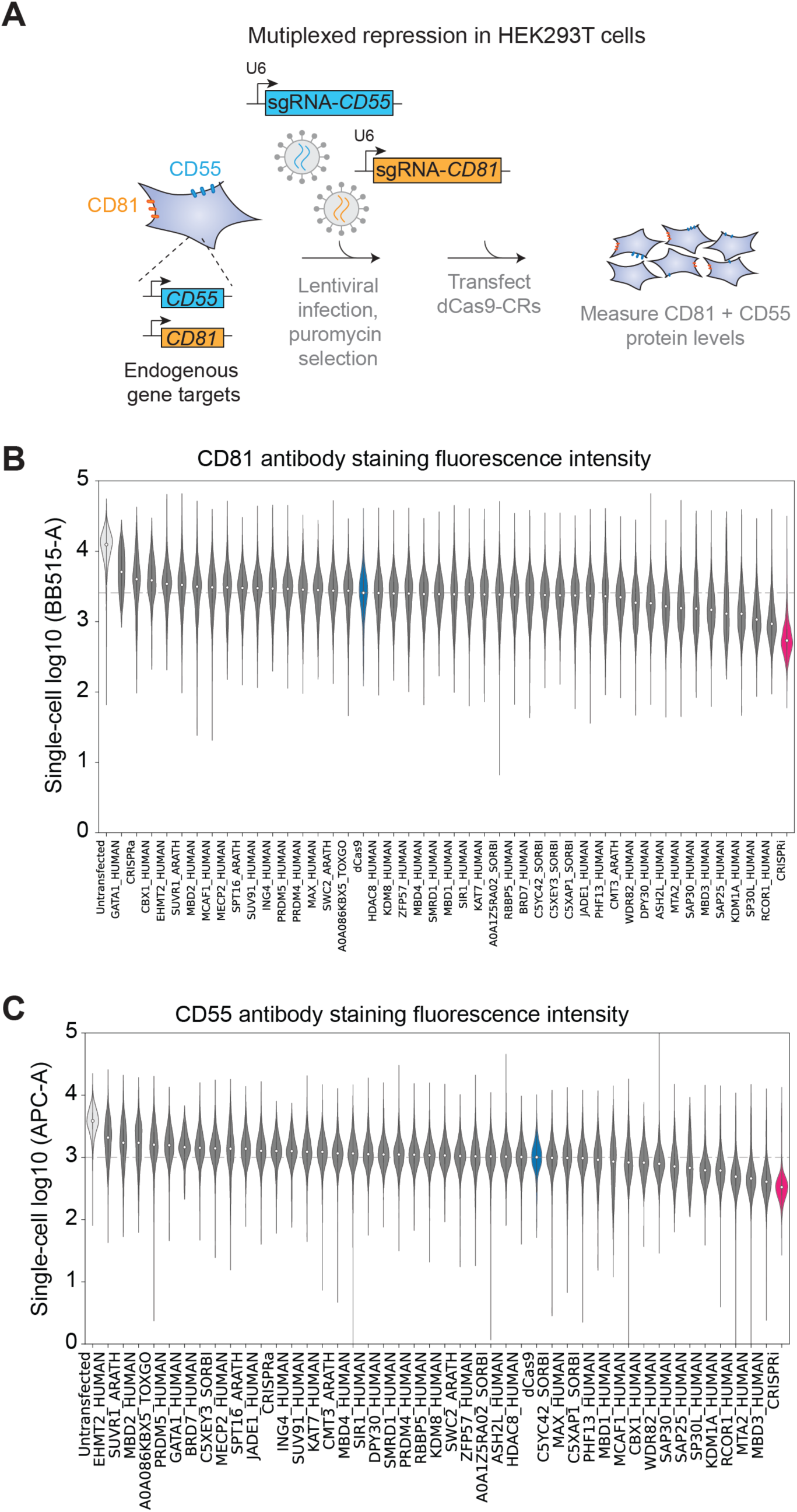
Multiplex assessment of CRs repression across endogenous gene targets. **(A)** Multiplex reporter assay for evaluating CR repression across endogenous targets *CD55* and *CD81*. **(B)** Violin plots showing single-cell antibody fluorescence intensity for all CRs at *CD55*. **(C)** same as in **(B)** but for *CD81*.

**Figure S3.**
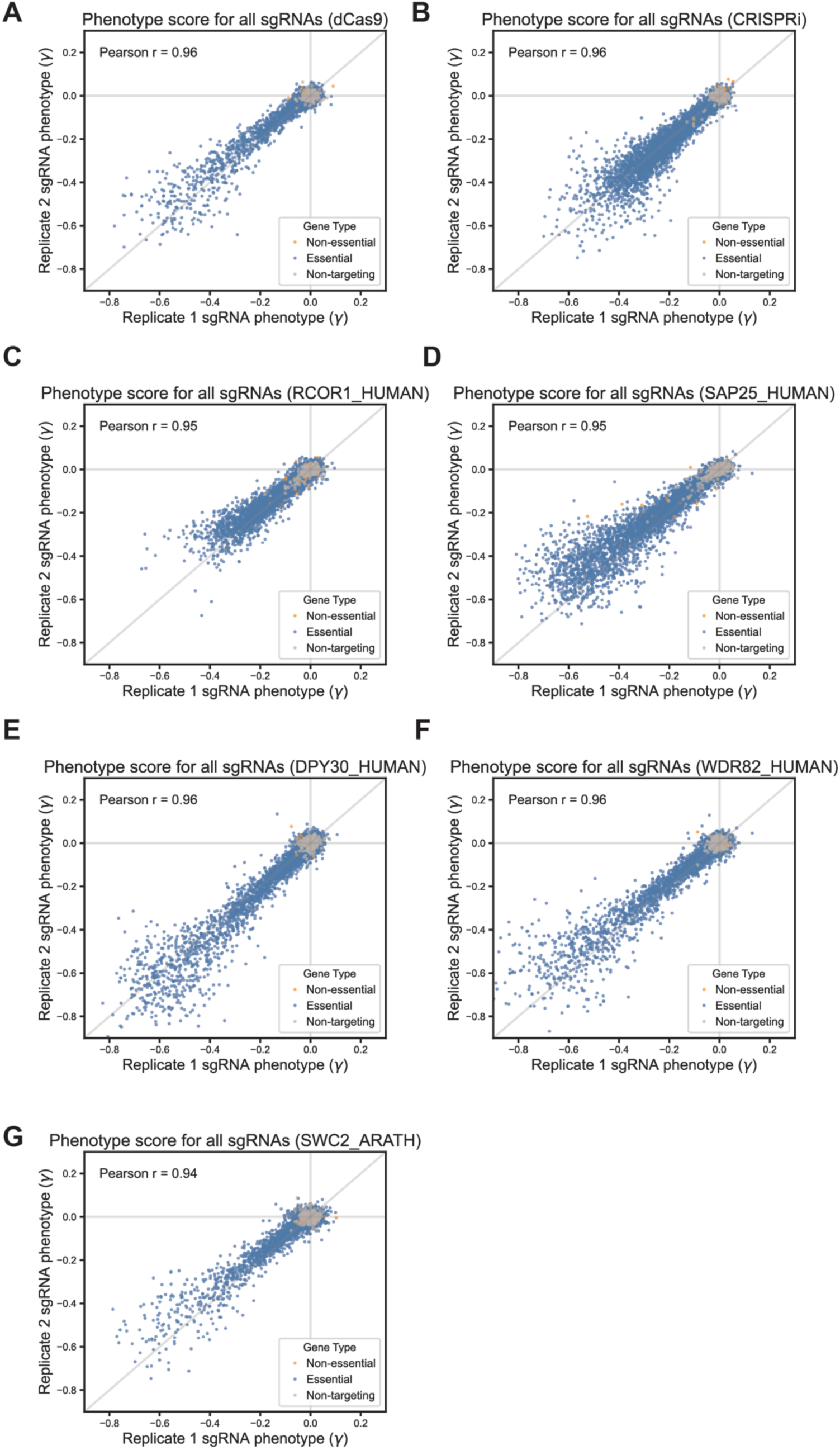
Replicate reproducibility of essential gene screen with CRs in K-562 cells. **(A)** Scatterplots showing replicate correlations of sgRNA phenotype scores (γ) from pooled essential gene screens performed in K-562 cells expressing dCas9 only. Each point represents an individual sgRNA targeting essential, non-essential, or non-targeting control genes. Pearson correlation coefficients (r) are indicated on each plot. **(B)** Same as **(A)** but for cells expressing CRISPRi (dCas9-KRAB). **(C)** Same as **(A)** but for cells expressing RCOR1_HUMAN (dCas9-RCOR1). **(D)** Same as **(A)** but for cells expressing SAP25_HUMAN (dCas9-SAP25). **(E)** Same as **(A)** but for cells expressing DPY30_HUMAN (dCas9-DPY30). **(F)** Same as **(A)** but for cells expressing WDR82_HUMAN (dCas9-WDR82). **(G)** Same as **(A)** but for cells expressing SWC2_ARATH (dCas9-SWC2).

**Figure S4.**
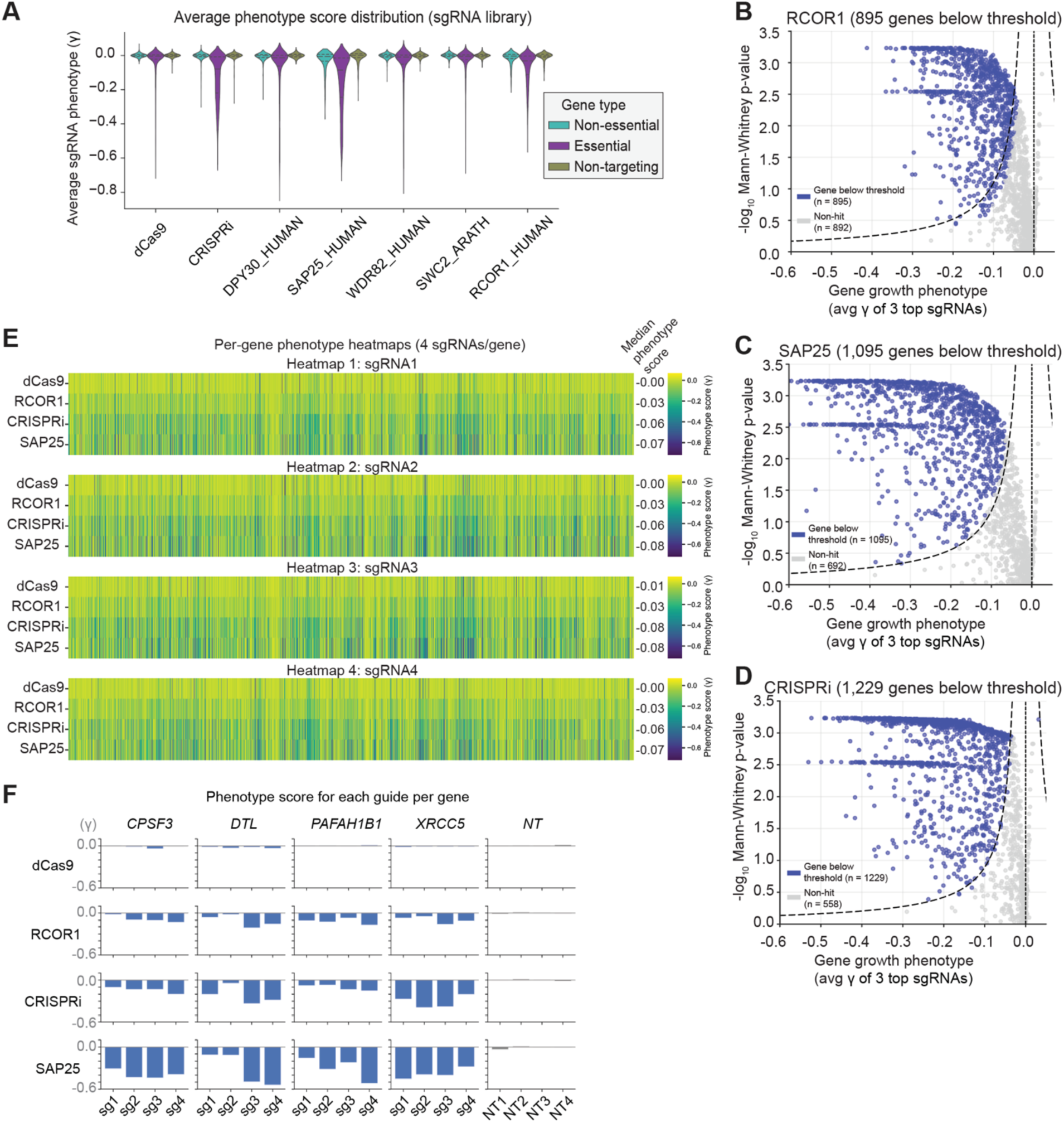
Tuning of cellular phenotypes across sgRNAs within the essential gene library. **(A)** Violin plots showing distribution of sgRNA phenotype scores across CRs and gene type (non-essential, essential, and non-targeting). **(B-D)** Volcano plots comparing the average gene growth phenotype of 3 top sgRNAs (x axis) and the –log10 Mann-Whitney p-value for RCOR1_HUMAN, SAP25_HUMAN and CRISPRi. Blue dots represent genes passing the threshold of significance and gray dots represent non-gene hits. **(E)** Per-gene heatmaps showing sgRNA-level phenotype scores for four sgRNAs per gene across dCas9, RCOR1, CRISPRi and SAP25. **(F)** Per-gene sgRNA-level phenotypes for representative essential genes targeted by four independent sgRNAs across effectors. Each bar represents the average phenotype score (γ) for individual sgRNAs targeting the indicated gene.

**Figure S5.**
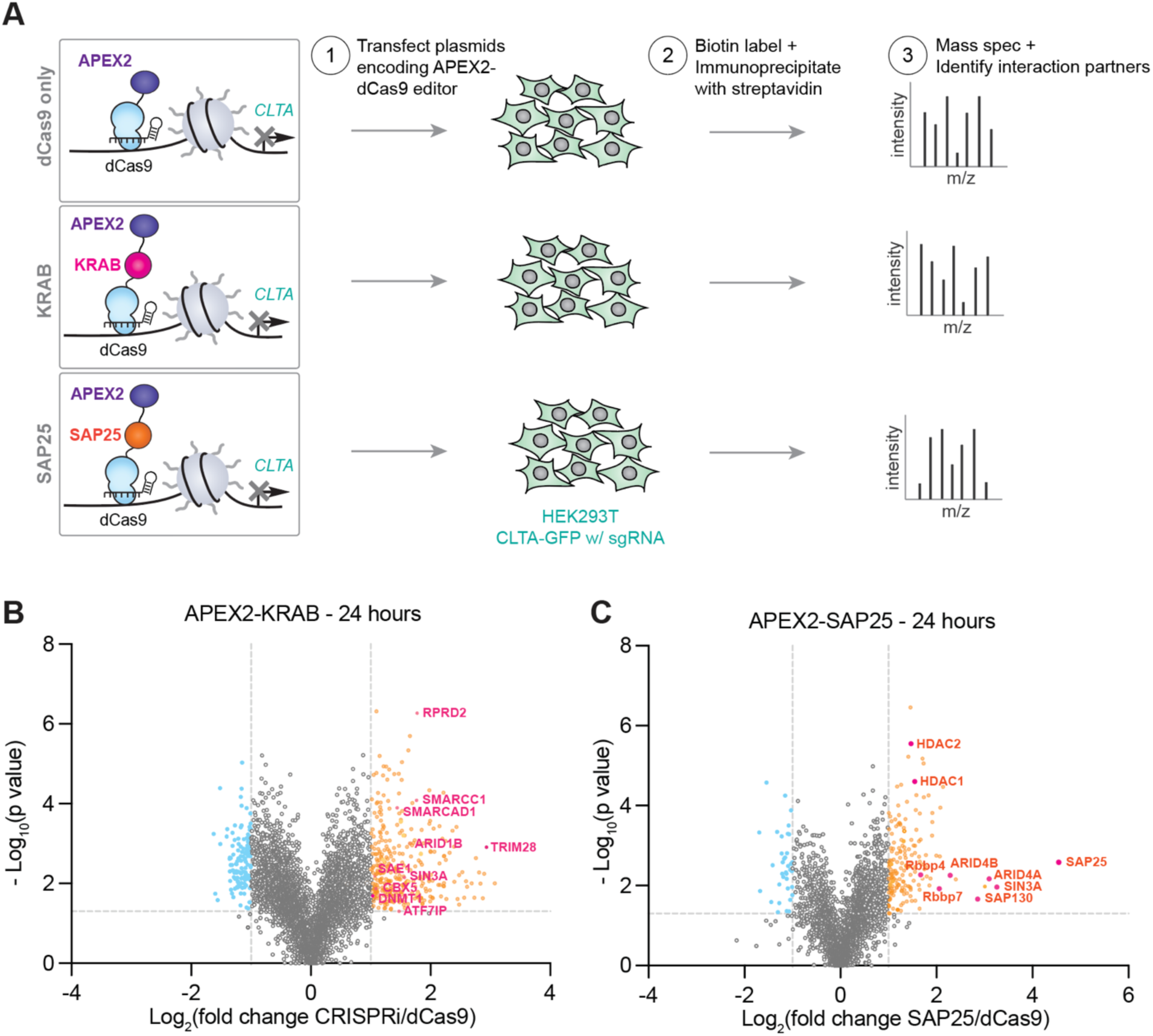
Proximity labeling identifies chromatin interaction partners of CRISPRi and SAP25 in HEK293T cells. **(A)** Schematic of the APEX2-based proximity labeling workflow used to map protein interactions of dCas9-fused repressors. HEK293T cells expressing dCas9-APEX2, dCas9-KRAB-APEX2, or dCas9-SAP25-APEX2 with sgRNAs targeting CLTA-GFP were transfected and labeled with biotin-phenol. Biotinylated proteins were enriched via streptavidin pulldown and analyzed by mass spectrometry to identify proximal interactors. **(B)** Volcano plots showing proteins significantly enriched in HEK293T cells expressing CRISPRi (APEX2-KRAB-dCas9) compared to APEX2-dCas9 control after 24 hours of labeling. **(C)** Same as in **(B)** but APEX2-SAP25-dCas9.

